# Loss of p21-activated kinase 4 (PAK4) suppresses pancreatic tumor progression and metastasis through regulating E-cadherin

**DOI:** 10.1101/2024.05.22.594599

**Authors:** Kui-Jin Kim, Milang Nam, Hee Young Na, Ji Hea Sung, Bo-Ram Park, Sung-Hyun Hwang, Woochan Park, Jeongmin Seo, Minsu Kang, Eun Hee Jung, Sang-A Kim, Koung Jin Suh, Ji Yun Lee, Ji-Won Kim, Se Hyun Kim, Jeong-Ok Lee, Yu Jung Kim, Keun-Wook Lee, Jee Hyun Kim, Soo-Mee Bang, Zev A. Wainberg, Jin Won Kim

## Abstract

Pancreatic ductal adenocarcinoma (PDAC) is characterized by a poor prognosis with early and frequent metastasis. While p21-activated kinase 4 (PAK4) has been implicated in cell migration, and invasion, the molecular mechanisms in PDAC remain unknown. In this study, we found that PAK4 overexpression was correlated with poor survival in PDAC patients through analysis of TCGA data. PAK4-amplified PDAC cells showed enhanced mobility in contrast with wild-type. PAK4 knockdown in PAK4 amplified cells inhibited cell migration, invasion, and displacement by increased and stabilized E-cadherin, which was attributed to decreased activity of Cdc42. PAK4 knock-in in PAK4 wild-type models enhanced cell migration, invasion, and displacement by reduced E-cadherin through elevated Cdc42 activity. PAK4 bounded to E-cadherin, Cdc42, and p120ctn in immunoprecipitation. In confocal imaging, the colocalization of PAK4, E-cadherin, p120ctn, and Cdc42 was also identified. In an orthotopic PDAC mouse model, PAK4 knockdown decreased primary tumor size and occurrence of malignant ascites by activation of E-cadherin. Notably, in patients’ tissue specimens, inverse correlation on expression of PAK4 and E-cadherin were also shown. In conclusion, our study highlights that PAK4 promotes invasive and metastatic behavior by regulating E-cadherin in PDAC. PAK4 could be a potential therapeutic target for PDAC patients.

**Graphical abstract:** 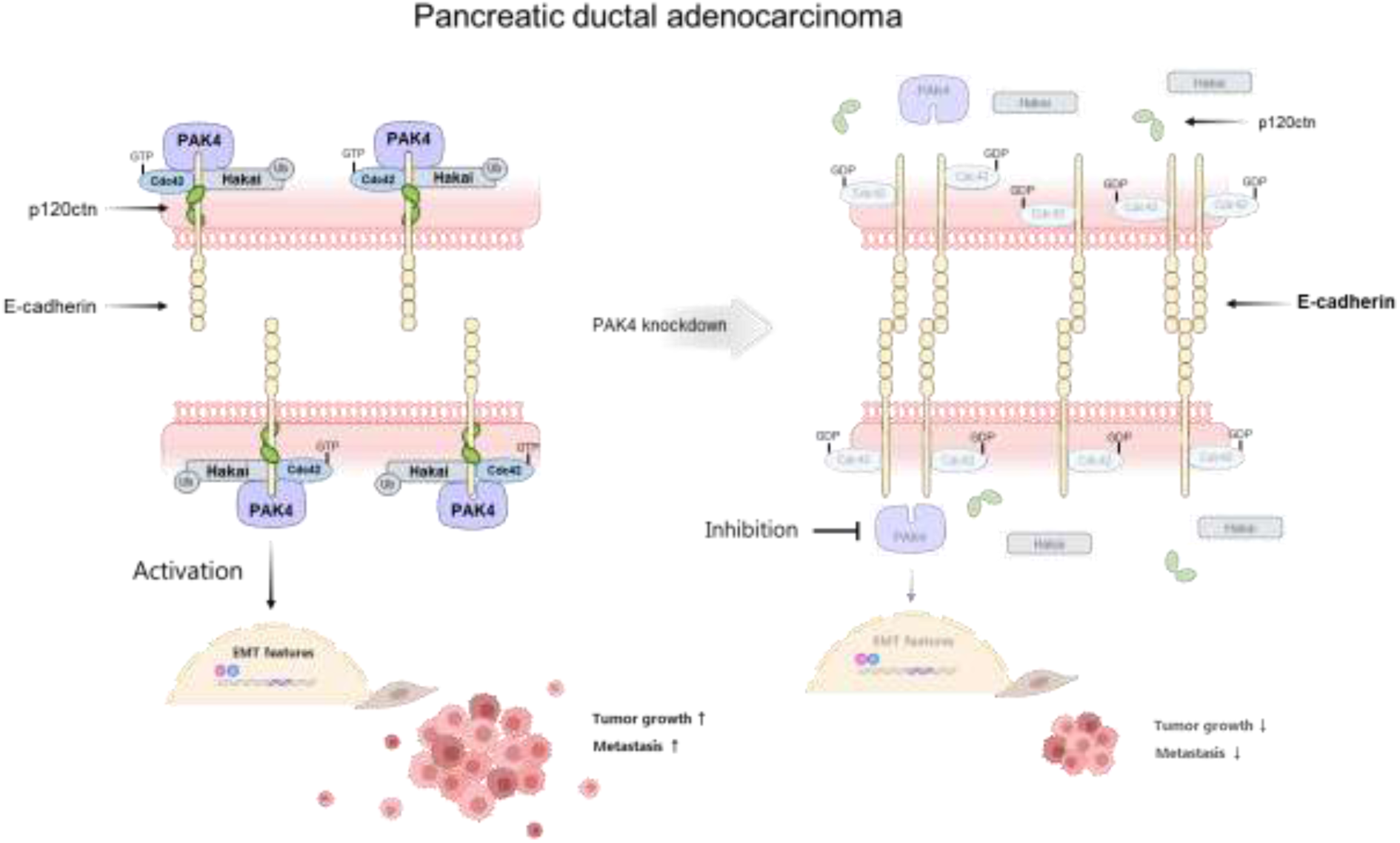

## Introduction

Pancreatic ductal adenocarcinoma (PDAC) arises from the pancreatic duct and is a highly aggressive and lethal form of cancer. It is characterized by early and frequent metastasis, advanced disease at diagnosis, a high recurrence rate after curative surgery, and a low treatment response, resulting in poor prognosis(*1–3*). Combined chemotherapies such as FOLFIRINOX and gemcitabine/nab-paclitaxel have improved the survival for those with metastatic disease and are also used as an adjuvant to surgical resection(*4–7*). Still, the prognosis for these patients remains poor(*2, 3, 8*). Despite increased knowledge about PDAC’s genetic and molecular characteristics, recent clinical trials evaluating targeted and immunotherapy approaches have demonstrated limited clinical efficacy(*2, 9*). Even in patients with high microsatellite instability (MSI-H) PDAC, the clinical efficacy of immunotherapy is lower than in other MSI-H tumors(*10*). Therefore, there is a critical unmet need for improved therapies for patients with PDAC.

The p21-activated kinase 4 (PAK4) is a member of the serine/threonine protein kinase family and has been implicated in the development and progression of various types of cancer, including PDAC(*11*). Previous studies revealed that PAK4 is overexpressed in PDAC, and its expression is strongly linked to disease progression and poor outcomes(*12*). The aberrant activation of PAK4 enhances tumor cell proliferation, migration, and invasion, while also promoting the epithelial-mesenchymal transition (EMT)(*13, 14*). This feature enables cancer cells to become more invasive, metastatic, and resistant to chemotherapy. In addition, PAK4 plays a crucial role in signaling pathways that regulate angiogenesis and the immune response(*15, 16*), both of which are critical for the growth and metastatic spread of melanoma and prostate cancer. Given its pivotal role in various cancer processes, elucidating the detailed molecular mechanisms by which PAK4 influences PDAC may hold immense promise for the development of treatment options.

Herein, our aim is to unveil the molecular mechanisms of PAK4 in a preclinical model to deepen the understanding of its specific function in PDAC. Our data may offer novel insights into PAK4 targeted therapy for PDAC, potentially advancing therapeutic strategies, particularly for patients with PDAC harboring PAK4 overexpression.

## Results

### PAK4 overexpression is associated with poor survival outcomes in PDAC

To evaluate the clinical impact of PAK4 overexpression (amplification and mRNA high) on prognosis in PDAC patients, we obtained publicly available Pancreatic Adenocarcinoma (TCGA, Firehose Legacy) from cBioportal (https://www.cbioportal.org/). Of 149 patients, 19 patients (13%) had PAK4 overexpression. Patients with PAK4 alteration showed significantly poorer disease-free survival (DFS) and overall survival (OS) compared with patients without PAK4 alteration (median DFS: 7.56 months vs. 17.05 months; median OS: 15.05 months vs. 20.17 months, respectively) (Fig. 1A-1C).

**Fig. 1.**
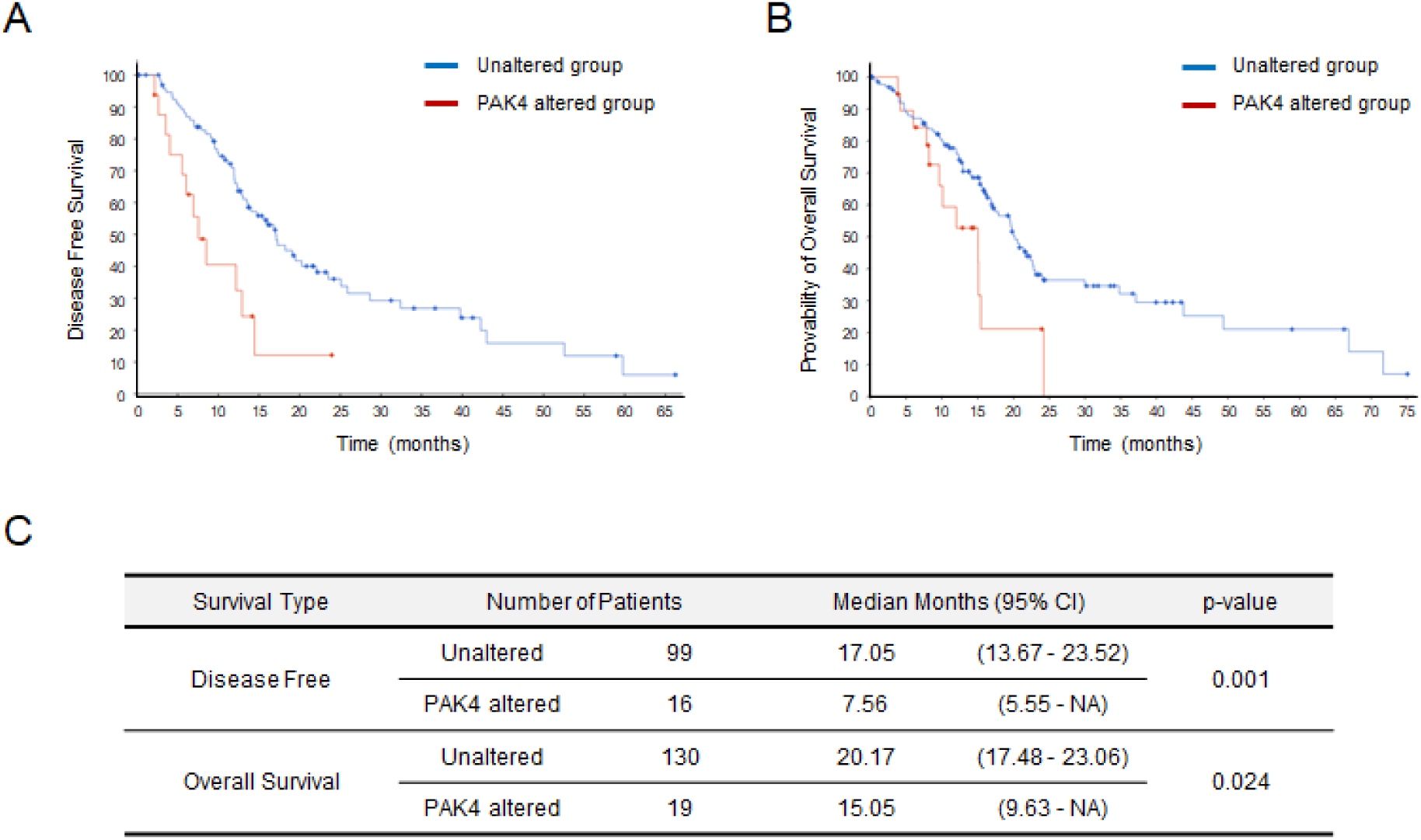
PAK4 alteration correlates with worse outcomes for PDAC patients. (A) Disease-free survival and (B) overall survival Kaplan–Meier curves were plotted according to PAK4 alteration, comparing patients with PAK4 alteration (red) and wild-type PAK4 (blue). (C) Analysis of the clinical features for disease-free survival and overall survival.

### PAK4 overexpression enhances cell migration in PDAC cell lines

Prior to exploring the function of PAK4 in cell proliferation and migration within PDAC cells, we first assessed the basal expression of PAK4 in a panel of PDAC cell lines by Western blot analysis. In PDAC cells harboring PAK4 genetic amplification (SU.86.86, PANC-1, PaTu-8988S, and PaTu-8988T), we observed a higher expression of PAK4 protein compared with wild-type cell lines (SNU-410, SNU-213, Capan-2, Capan-1, AsPC-1, and MIAPaCa-2), as shown in Figure 2A and 2B. To elucidate the role of PAK4 overexpression in PDAC cells, we conducted the cell proliferation assays. As shown in Figure 2C, it was observed the there is no correlation with PAK4 overexpression. We next analyzed the migration of PDAC cells harboring PAK4 overexpression and PAK4 wild-type cells using a Boyden chamber migration assay, a commonly used method to measure the migratory potential of cells in vitro(*17*). As shown in Figure 2D and 2E, PDAC cells harboring PAK4 overexpression (SU.86.86, PANC-1, and PaTu-8988T) exhibited increased migration activity compared with wild-type cells. This effect was independent of cell proliferation, indicating that PAK4 overexpression may stimulate cell migration in a cell proliferation-independent manner..

**Fig. 2.**
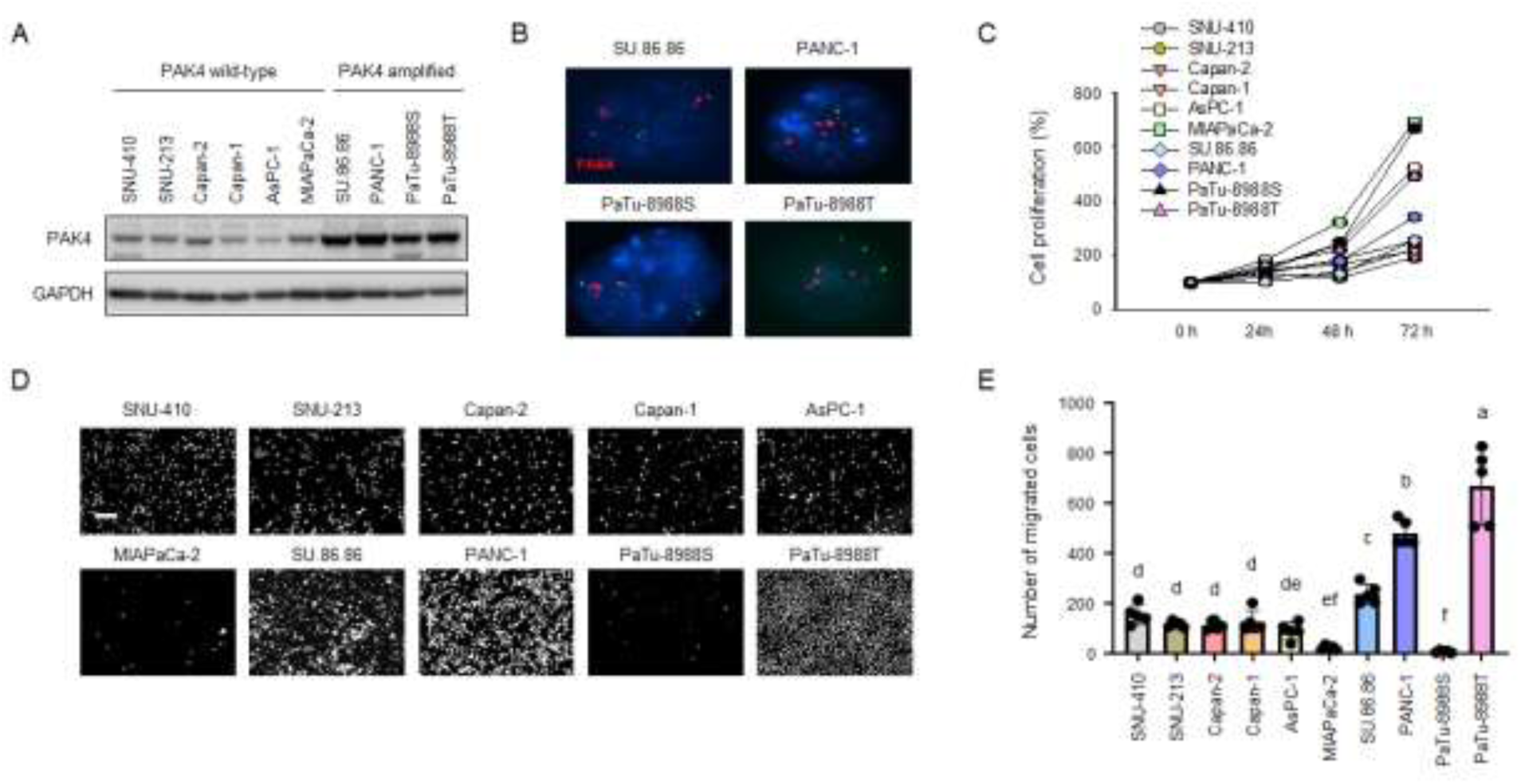
PAK4 amplification enhances cell migration in PDAC cells in a cell proliferation-independent manner. (A) The protein expression levels of PAK4 were evaluated in SNU-410, SNU-213, Capan-2, Capan-1, AsPC-1, MIAPaCa-2, SU.86.86, PANC-1, PaTu-8988s, and PaTu-8988T cells. GAPDH was used as a protein loading control. (B) Fluorescence in situ hybridization (FISH) was performed using a probe within the amplified genomic region labeled with Cy3 (red) and a centromere reference probe labeled with FITC (Green). (C) Cell proliferation assays with PAK4 wild-type cells (SNU-410, SNU-213, Capan-2, Capan-1, AsPC-1, and MIAPaCa-2) and PAK4 amplified cells (SU.86.86, PANC-1, PaTu-8988s, and PaTu-8988T) were performed using a CellTiter-Glo assay. Data are presented as mean ± SD. (D) The indicated PDAC cells were allowed to migrate for 20 hours. The experiment was performed in triplicate. (E) Data are analyzed by ImageJ software and presented as mean ± SD.

### Loss of PAK4 reduces cell migration in PAK4 overexpression PDAC cells

To further explore the impact of PAK4 on cell migration, we knocked down PAK4 in the PAK4 over-expresed cell lines (SU.86.86, PANC-1, and PaTu-8988T) using PAK4 shRNA expressed from a pLKO.1 puro plasmid (Fig. 3A). PAK4 knockdown using two independent shRNAs significantly reduced single-cell migration of SU.86.86, PANC-1, and PaTu-8988T cells (Fig. 3B). A comparable effect was seen for collective cell migration (Fig. 3C). In addition, we found that reduction of PAK4 levels in SU.86.86 and PANC-1 cells significantly inhibited cell invasion when compared with control cells (Fig. 3D). These results indicate that reducing levels of PAK4 disrupts single cell migration and collective cell migration in PDAC cells harboring PAK4 overexpression, potentially by modulating cell mobility.

**Fig. 3.**
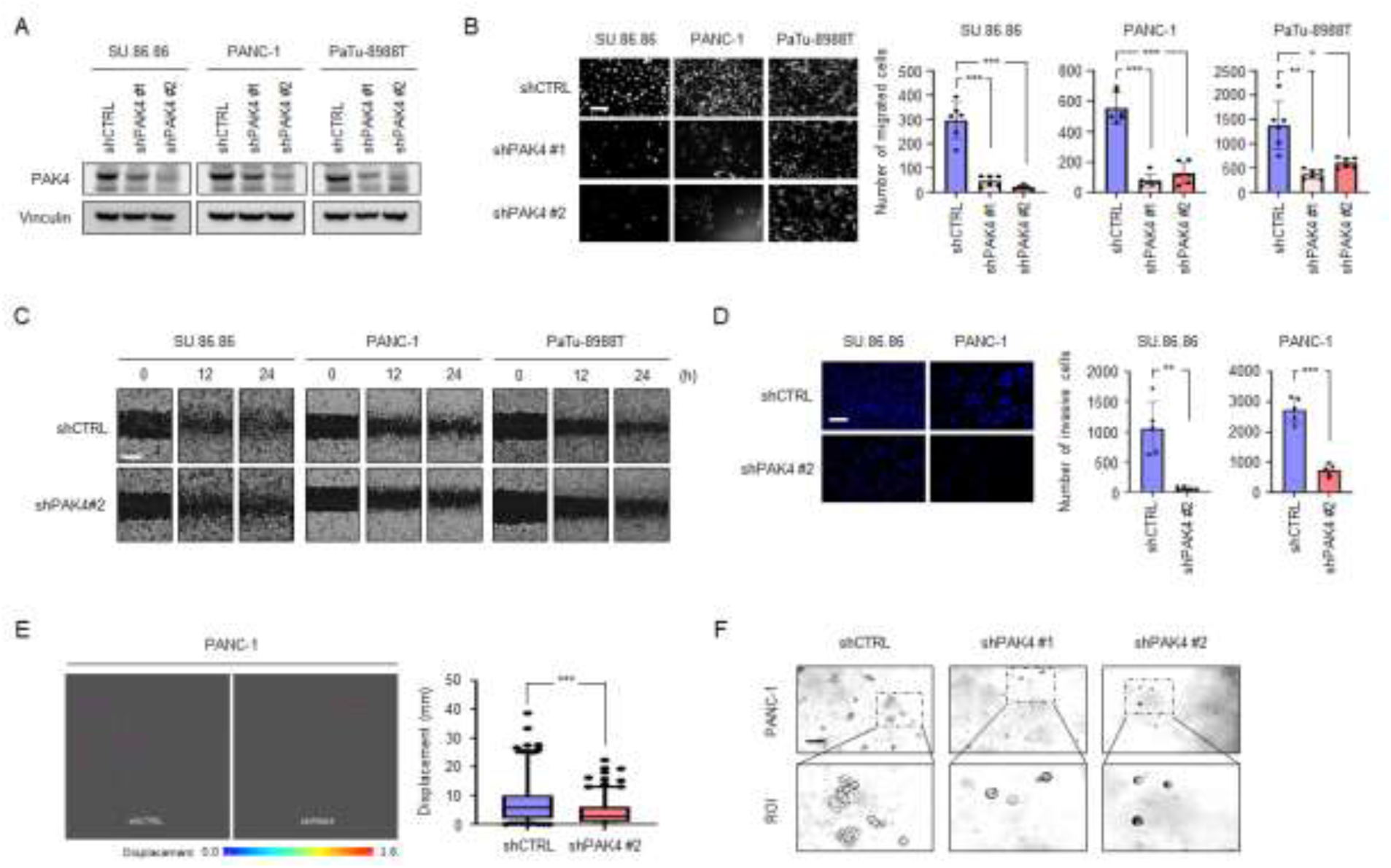
PAK4 knockdown suppresses cell migration, invasion, displacement, and sphere-forming ability in PDAC cells with PAK4 amplification. (A) The protein expression levels of PAK4 were evaluated in SU.86.86, PANC-1, and PaTu-8988T cells with control shRNA or PAK4 knockdown. Vinculin was used as a protein-loading control. (B) A single-cell migration assay was used to evaluate the effect of PAK4 knockdown on the migration of SU.86.86, PANC-1, and PaTu-8988T cells. Scale bar, 100 μm (C) Cell migration was evaluated using a wound-healing assay. (D) A Matrigel-coated transwell assay was used to evaluate the effect of PAK4 knockdown on the invasive potential of SU.86.86 and PANC-1 cells. Scale bar, 100 μm (E) Cell displacement was analyzed by IncuCyte imaging in PANC-1 cells with shCTRL or shPAK4. (F) The sphere-forming potency of PANC-1 cells with PAK4 knockdown. Scale bar, 50 μm

To test whether PAK4 knockdown could affect cell mobility, we examined the effect of PAK4 knockdown on cell displacement *in vitro*. As shown in Figure 3E, PAK4 knockdown in PANC-1 cells significantly decreased cell displacement. In a sphere formation assay, knocking down PAK4 in PANC-1 cells caused a dramatic change in sphere morphology. While control cells formed spheres with irregular budding, PAK4-depleted spheres were smooth and round, further highlighting the enhanced cell migration and invasion capacity of cells with PAK4 amplification (Fig. 3F). These data suggest that inhibition of PAK4 expression may functionally disrupt the migratory potential of PAK4 amplified cells.

### PAK4 knockdown upregulates E-cadherin in PDAC cells

We next analyzed signaling pathways related to EMT to clarify the potential molecular mechanisms by which PAK4 knockdown affects migration, invasion, and displacement in PDAC cells. In both SU.86.86 and PANC-1 PAK4 amplified cells,, knocking down PAK4 significantly elevated E-cadherin mRNA expression compared to controls, while it significantly reduced the expression of N-cadherin, ZEB1, ZEB2, Snail, and Vimentin (Fig. 4A). PAK4 knockdown also significantly decreased the expression of Slug in SU.86.86 cells, a finding not not apparent in PANC-1 cells. We also observed consistent changes in protein levels upon reduction of PAK4 in both SU.86.86 and PANC-1 cell lines, including increased E-cadherin and modulation of other EMT-associated factors,(p-p120ctn, ZEB1, Snail, Slug, and Vimentin) (Fig.4B). Coversely, knockdown of PAK4 elevated the expression of Zonula occludens-1 (ZO-1) in SU.86.86 cells and CLDN-1 in PANC-1 cells.

**Fig. 4.**
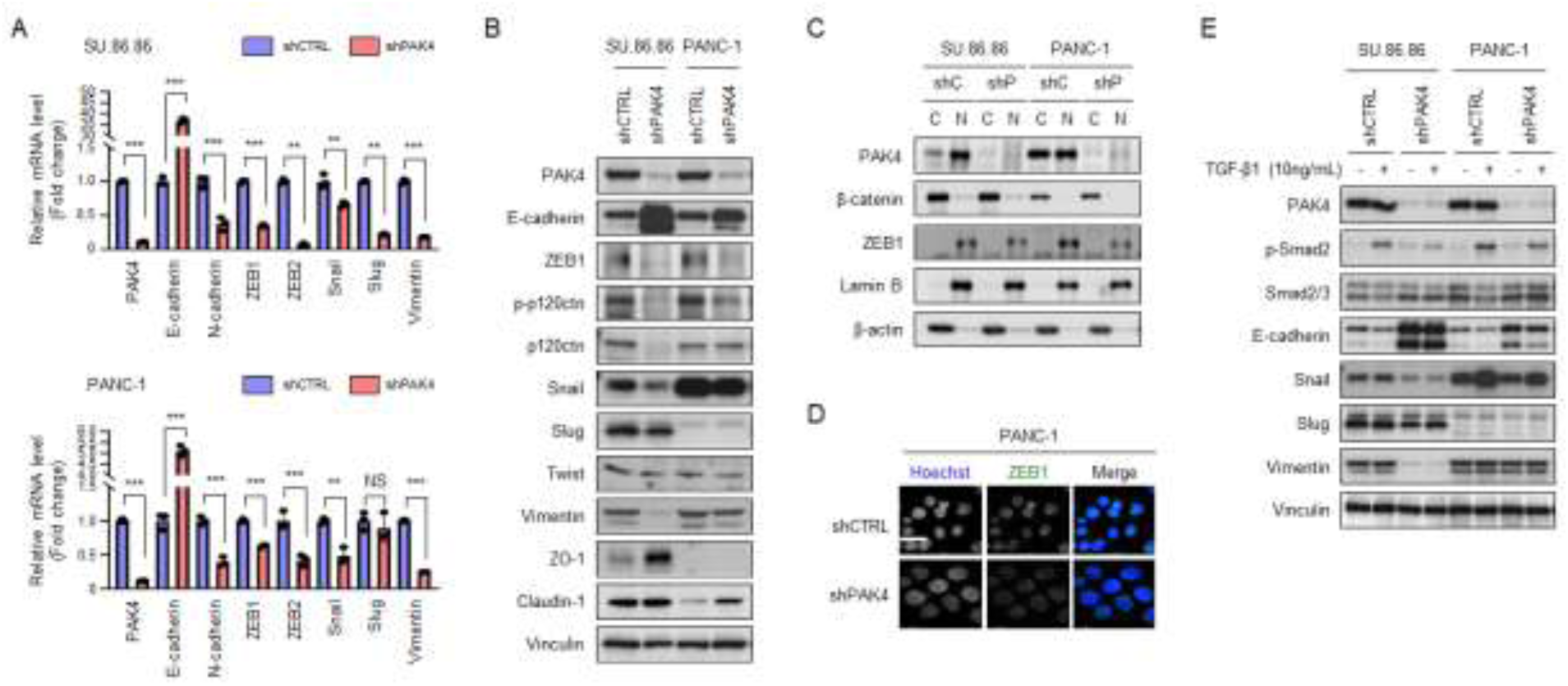
PAK4 knockdown increases the epithelial markers and decreases the mesenchymal markers in PDAC cells harboring PAK4 amplification. (A) Real-time PCR analysis was performed on total RNA to evaluate mRNA levels of PAK4 and its target genes as indicated in SU.86.86 and PANC-1 cells stably expressing shCTRL or shPAK4. Vinculin was used as a protein-loading control. (B) SU.86.86 and PANC-1 cells stably expressing shCTRL or shPAK4 were subjected to Western blot analysis to detect the levels of EMT-associated factors. (C) The cytoplasmic and nuclear lysates of SU.86.86 and PANC-1 cells stably expressing shCTRL or shPAK4 were analyzed by Western blot with the indicated antibodies. (D) The intracellular expression ZEB1 was analyzed by immunofluorescence in PANC-1 cells, Scale bar, 20 μm. (E) SU.86.86 and PANC-1 cells were treated with 10ng/mL TGF-β1 for 20 hours, and then the expression of PAK4, p-Smad2, Smad2/3, and E-cadherin were analyzed by Western blot. Vinculin was used as a protein-loading control.

The nuclear co-localization of β-catenin and ZEB1 has been shown to promote an EMT phenotype and repress E-cadherin expression during tumor progression(*18–20*). To gain a more comprehensive understanding of how PAK4 regulates E-cadherin expression, we investigated whether PAK4 knockdown could influence the localization and expression of β-catenin and ZEB1 in SU.86.86 and PANC-1 cell lines, as shown in Figure 4C. Analysis of the cytoplasmic and nuclear fractions revealed that the majority of β-catenin was located in the cytoplasm and was not affected by PAK4 knockdown in SU.86.86 or PANC-1 cells. In contrast, the majority of ZEB1 was found in the nucleus, and its expression was slightly decreased upon PAK4 knockdown in both SU.86.86 and PANC-1 cells compared with their respective controls (Fig. 4D).

There is a well-established interaction between the TGF-β1 and SMAD signaling pathways, which together regulate the canonical EMT pathway(*21, 22*). To determine the involvement of PAK4 in the canonical TGF-β1-regulated EMT pathway, we treated PDAC cells with TGF-β1. As shown in Figure 4E, TGF-β1 increased the levels of p-SMAD2 in shCTRL SU.86.86 and PANC-1 cells at 24 hours by approximately 80% and 60%, respectively, and decreased the expression of E-cadherin by approximately 20% and 30%, respectively. In contrast, shPAK4 SU.86.86 and PANC-1 cells exhibited a blunted p-SMAD2 response to TGF-β1 treatment. In addition, PAK4 knockdown inhibited the TGF-β1-mediated suppression of E-cadherin in SU.86.86 and PANC-1 cells that was observed in control cells. These findings suggest that PAK4 knockdown may involve non-canonical regulation of E-cadherin expression in SU.86.86 and PANC-1 cells.

### The loss of PAK4 stabilizes E-cadherin expression

To determine the mechanism by which E-cadherin levels are stabilized upon PAK4 knockdown, we treated cells with the protein synthesis inhibitor, cycloheximide (CHX). E-cadherin was rapidly degraded in shCTRL PANC-1 cells, whereas its levels remained stable in shPAK4 PANC-1 cells (Fig. 5A). In contrast, CHX induced rapid destabilization of ZEB1, with a decrease in ZEB1 protein levels within 3 hours of CHX treatment. The half-life of ZEB1 was 6 hours in both shCTRL and shPAK4 PANC-1 cells.

**Fig. 5.**
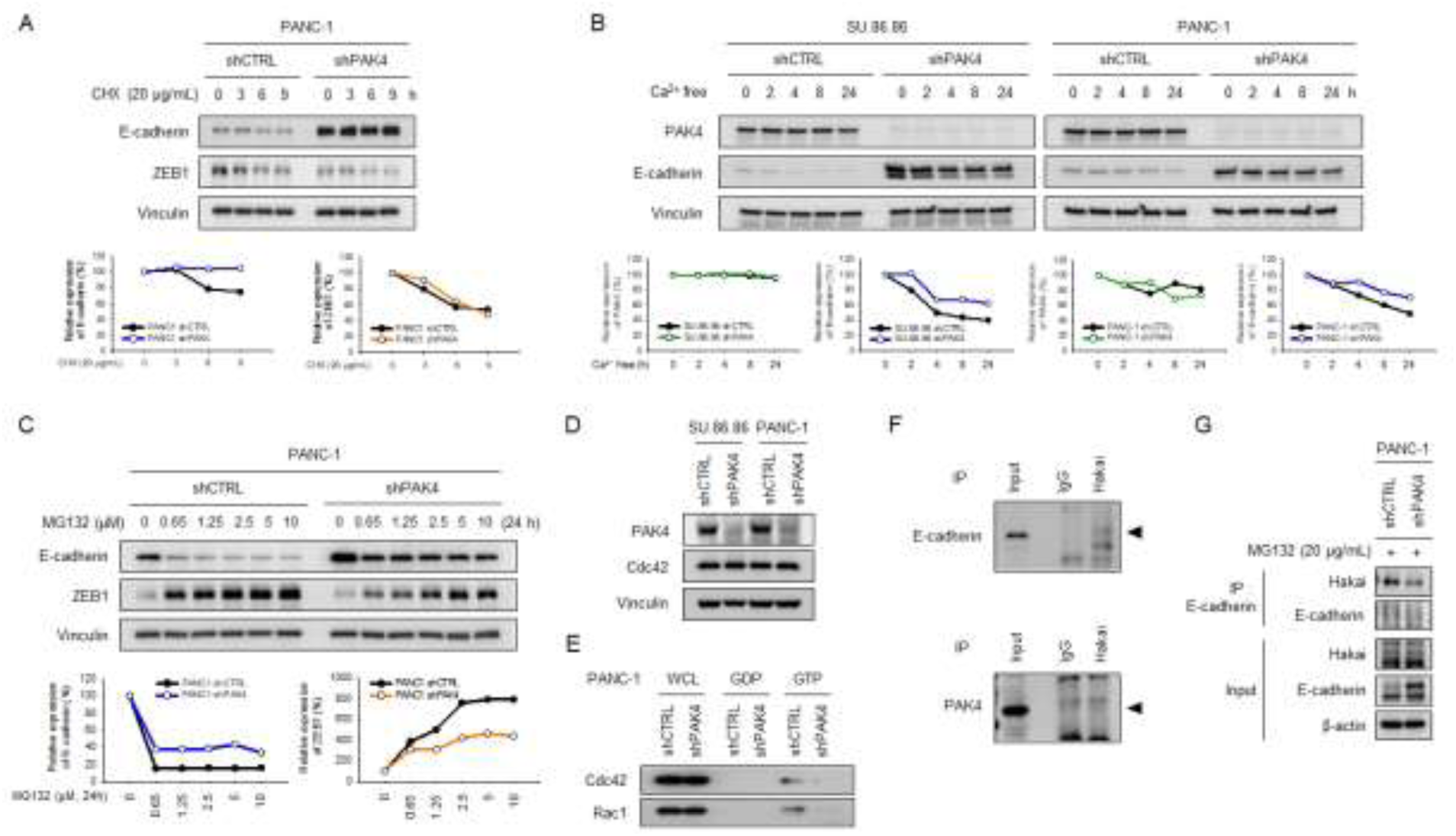
PAK4 stabilizes E-cadherin through the inhibition of Cdc42 activity. (A) Left panel: PANC-1 cells were treated with 20μg/mL cycloheximide (CHX), harvested at the indicated time points, and Western blotted with antibodies against E-cadherin, ZEB1, and Vinculin. Right panel: Quantification of E-cadherin and ZEB1 protein levels (normalized to Vinculin). (B) Upper panel: SU.86.86 and PANC-1 cells were seeded overnight in 1%P/S/10%FBS/DMEM and then switched to DMEM without Ca^2+^ for the indicated times. E-cadherin, ZEB1, and Vinculin were detected by Western blot. Bottom panel: Quantification of E-cadherin and ZEB1 protein levels (normalized to Vinculin). (C) Upper panel: PANC-1 cells stably expressing shCTRL or shPAK4 were treated with the indicated concentration of MG132, harvested at 24 h, and Western blotted with antibodies against E-cadherin, ZEB1, and Vinculin. Bottom panel: Quantification of E-cadherin and ZEB1 protein levels (normalized to Vinculin). (D) PANC-1 cells stably expressing shCTRL or shPAK4 were incubated in serum-free medium for the indicated times. Activated Cdc42 and Rac1 were affinity-precipitated from whole cell lysate by incubating with GST-PBD-agarose. Bound Cdc42 and Rac1 were detected by western blot for Cdc42 and Rac1. (E) SU.86.86 and PANC-1 cells were subjected to Western blot analysis to detect the levels of PAK4 and Cdc42. Vinculin was used as a protein-loading control. (F) Interaction between endogenous PAK4 and endogenous Hakai in SU.86.86 cells and interaction between endogenous E-cadherin and endogenous Hakai in SU.86.86 cells. Cells were treated with serum-free medium for 24 h, and immunoprecipitation with anti-Hakai was performed before Western blotting with the indicated antibodies. (G) Cell lysates from PANC-1 cells were immunoprecipitated with antibodies against E-cadherin. Proteins immunoprecipitated by E-cadherin were then Western blotted for endogenous Hakai and E-cadherin. Whole cell lysate from PANC-1 was Western blotted with the indicated antibodies.

E-cadherin lysosomal degradation occurs in response to Ca^2+^ depletion in a Cdc42-dependent manner(*23*). To gain deeper insight into how PAK4 knockdown influences the regulation of E-cadherin, we examined the impact of Ca^2+^ depletion on E-cadherin levels. The expression of PAK4 remained unaffected by the depletion of Ca^2+^ in both shCTRL and shPAK4 SU.86.86 and PANC-1 cells. Notably, the withdrawal of Ca^2+^ led to a gradual decrease in E-cadherin expression levels in SU.86.86 and PANC-1 cells transfected with shCTRL. This decline commenced within 2 hours and reached a minimum by 24 hours (Fig. 5B). In contrast, the depletion of Ca^2+^ had a lesser effect on the expression level of E-cadherin in PAK4 knockdown cells. We note that in shCTRL cells, proteasomal inhibition with MG132 led to the stabilization of ZEB1 and a concomitant reduction in E-cadherin protein levels (Fig. 5C). On the other hand, PAK4 knockdown inhibited this ZEB1-associated reduction in E-cadherin protein when PANC-1 cells were treated with MG132. Taken together, these data suggest that the presence of PAK4 decreases the stability of E-cadherin in a ZEB1-independent manner.

Next, we analyzed the expression and activity of Cdc42 in SU.86.86 and PANC-1 cells to elucidate the relationship between PAK4 and E-cadherin (Fig. 5D and 5E). We found similar levels of total protein but reduced levels of activated Cdc42 in shPAK4 cells compared to the control. These results suggest that PAK4 is involved in the activation of Cdc42, but not in the expression of Cdc42. Thus, the loss of PAK4 leads to reduced activation of Cdc42, which is linked to the repression of E-cadherin degradation.

E-cadherin degradation is a complex process that is regulated by various factors, including the activity of E3 ubiquitin ligases like Hakai(*24*). To understand how PAK4 regulates E-cadherin stability, we performed immunoprecipitation assays to examine the interaction between E-cadherin and Hakai in PANC-1 cells. As shown in Figure 5F, immunoprecipitation revealed that Hakai binds to E-cadherin, as well as PAK4. In addition, E-cadherin exhibited less binding with Hakai in shPAK4 PANC-1 cells compared to shCTRL PANC-1 cells (Fig. 5G), suggesting that PAK4 knockdown reduced the degradation of E-cadherin through the ubiquitin-proteasome system. These results suggest that PAK4 plays the key role in regulating the stability of E-cadherin, which is a critical protein for maintaining cell-cell adhesion and tissue organization.

### PAK4 binds to E-cadherin, p120ctn, and Cdc42 in SU.86.86 cells

The aforementioned results prompted us to hypothesize that PAK4 may interact with E-cadherin and Cdc42 to regulate cell migration and invasion. Whole cell lysates of SU.86.86 cells were subjected to anti-PAK4 and anti-Cdc42 immunoprecipitation. As shown in Figure 6A, PAK4 bound to E-cadherin and Cdc42. Additionally, p120ctn, which regulates E-cadherin(*25*), was found in the anti-PAK4 immunoprecipitation. We also observed that Cdc42 interacted with PAK4, E-cadherin, and p120ctn. Furthermore, confocal imaging revealed the colocalization of PAK4, E-cadherin, p120ctn, and Cdc42 in SU.86.86 cells, as shown in Figure 6B-6G. Collectively, these results suggest that PAK4 inhibits the stabilization of E-cadherin through regulation of non-canonical EMT signaling in PDAC cells harboring PAK4 overexpression.

**Fig. 6.**
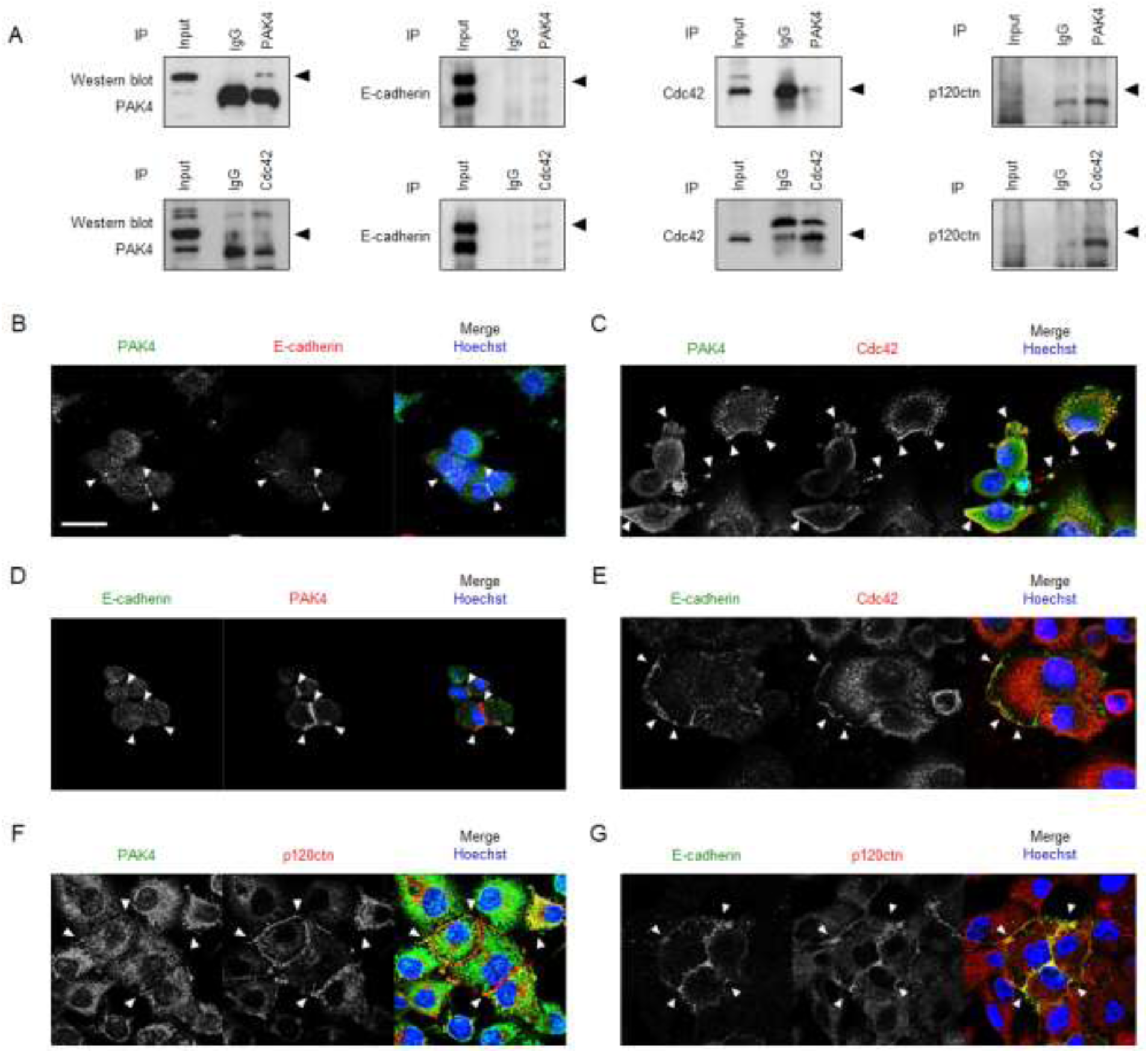
Endogenous PAK4 interacts with E-cadherin, p120ctn, and Cdc42 in PDAC cells. (A) Cell lysates from SU.86.86 cells were subjected to immunoprecipitation with antibodies against PAK4, Cdc42, or a control IgG antibody. Proteins immunoprecipitated by PAK4 or Cdc42 were then Western blotted for endogenous PAK4, E-cadherin, Cdc42, and p120ctn. (B) PAK4 (green) and E-cadherin (red) colocalize (yellow in merged) in SU.86.86 cells. A triple merge of PAK4, E-cadherin, and Hoechst is shown. Size bars correspond to 20 μm. Arrowheads indicate prominent sites of colocalization. (C) PAK4 (green) and Cdc42 (red) colocalize (yellow in merged) in SU.86.86 cells. (D) E-cadherin (green) and PAK4 (red) colocalize (yellow in merged) in SU.86.86 cells. (E) E-cadherin (green) and Cdc42 (red) colocalize (yellow in merged) in SU.86.86 cells. (F) PAK4 (green) and p120ctn (red) colocalize (yellow in merged) in SU.86.86 cells. (G) E-cadherin (green) and p120ctn (red) colocalize (yellow in merged) in SU.86.86 cells.

### Gain of PAK4 expression enhances cell migration and displacement by suppressing E-cadherin and elevating Cdc42 activity in PAK4 wild-type cells

We tested whether overexpression of PAK4 could stimulate cell migratory potential and mobility through modulation of E-cadherin expression. Overexpression of PAK4 in 293T and AsPC-1 cells, which do not exhibit PAK4 overexpression, dramatically decreased E-cadherin expression compared with control 293T and AsPC-1 cells (Fig. 7A). In addition, overexpression of PAK4 significantly boosted the migration and invasiveness of AsPC-1 cells (Fig. 7B and 7C), and led to a noticeable increase in cell displacement (Fig. 7D). Finally, overexpression of PAK4 resulted in an increase in the levels of activated Cdc42 in AsPC-1 cells (Fig. 7E).

**Fig. 7.**
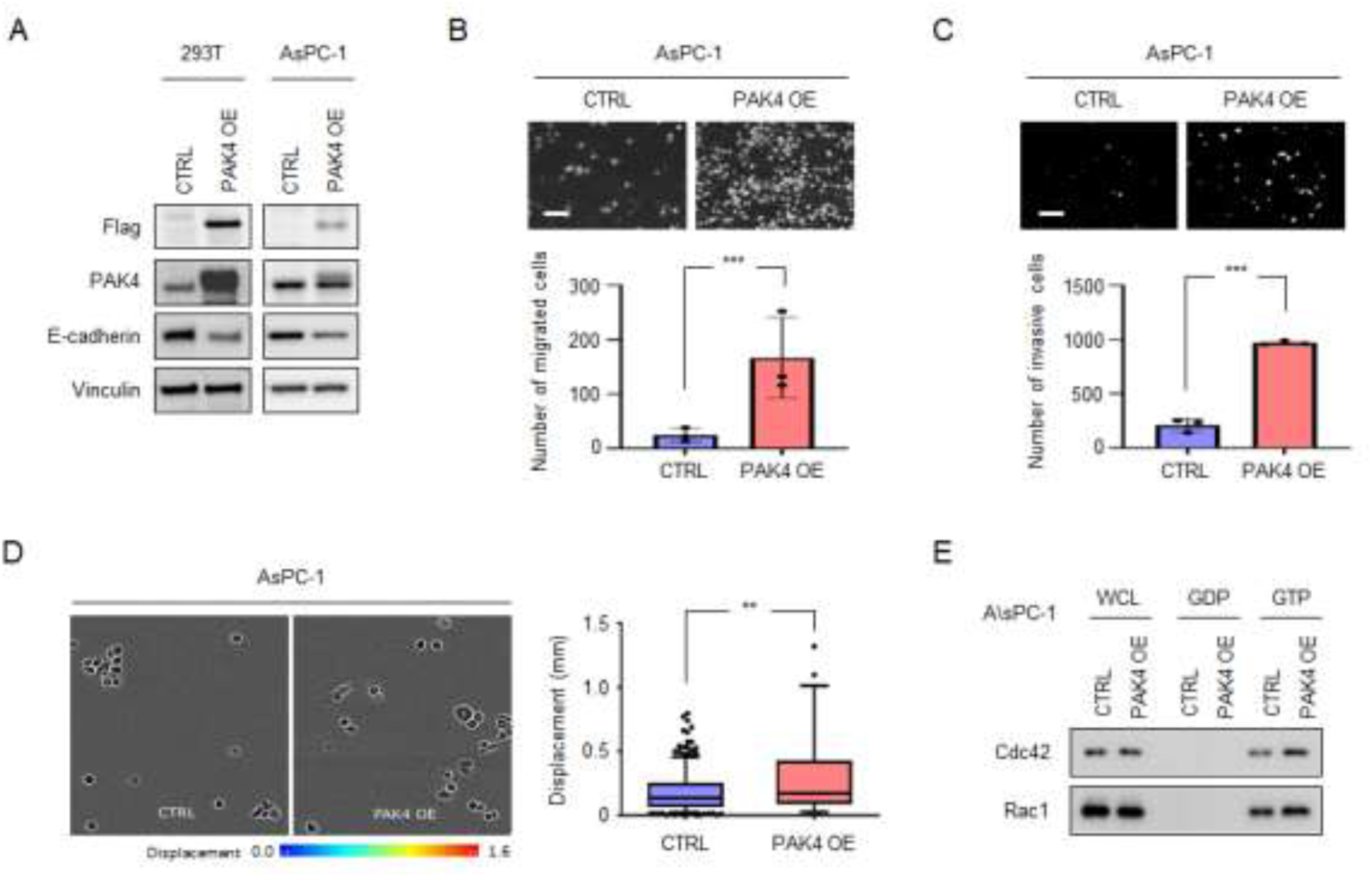
Overexpression of PAK4 increases cell migration, invasion, and displacement in 293T and AsPC-1 cells harboring PAK4 wild-type. (A) The protein expression levels of PAK4 and E-cadherin were evaluated in control 293T and AsPC-1 cells and those transfected with a PAK4 overexpression plasmid. Vinculin was used as a protein-loading control. (B) A single-cell migration assay was used to evaluate the effect of PAK4 overexpression on the migration of AsPC-1 cells. Scale bar, 100 μm (C) A Matrigel-coated transwell assay was used to evaluate the effect of PAK4 overexpression on the invasive potential of AsPC-cells. Scale bar, 100 μm (D) A cell displacement assay, analyzed by IncuCyte imaging, was used to evaluate the effect of PAK4 overexpression on AsPC-1 cells. (E) Control AsPC-1 cells or those with PAK4 overexpression were incubated in serum-free medium for the indicated times. Activated Cdc42 and Rac1 were affinity-precipitated from whole cell lysate by incubating with GST-PBD-agarose. Bound Cdc42 and Rac1 were detected by western blot for Cdc42 and Rac1.

### In a PDAC orthotopic model, loss of PAK4 inhibits primary tumor growth and metastasis

We investigated the effects of PAK4 knockdown on primary tumor growth and metastasis in a PDAC orthotopic mouse model. We transfected PANC-1 cells with lentiviruses that stably expressed either luciferase or the puromycin resistance gene, along with shRNAs targeting PAK4. We then transplanted these PANC-1 cells with and without PAK4 knockdown into the pancreases of the mice orthotopically. PAK4 knockdown had a negligible effect on body weight compared to the control (Fig. 8A), but significantly reduced cancer progression and primary tumor burden (Fig. 8B and 8C). Of note, approximately 66.7% ascites development was observed in mice transplanted with shCTRL PANC-1 cells (Fig. 8D). This phenomenon was not observed for mice with the PAK4 knockdown cells. A luciferase activity assay revealed the presence of luciferase-positive cells in the ascites of the mice transplanted with shCTRL PANC-1 cells, indicating peritoneal metastasis of a potential population of cancer cells with metastatic characteristics (Fig. 8E). PAP and H&E staining provided clearer visualization of these cancer cells in the ascites (Fig. 8F).

**Fig. 8.**
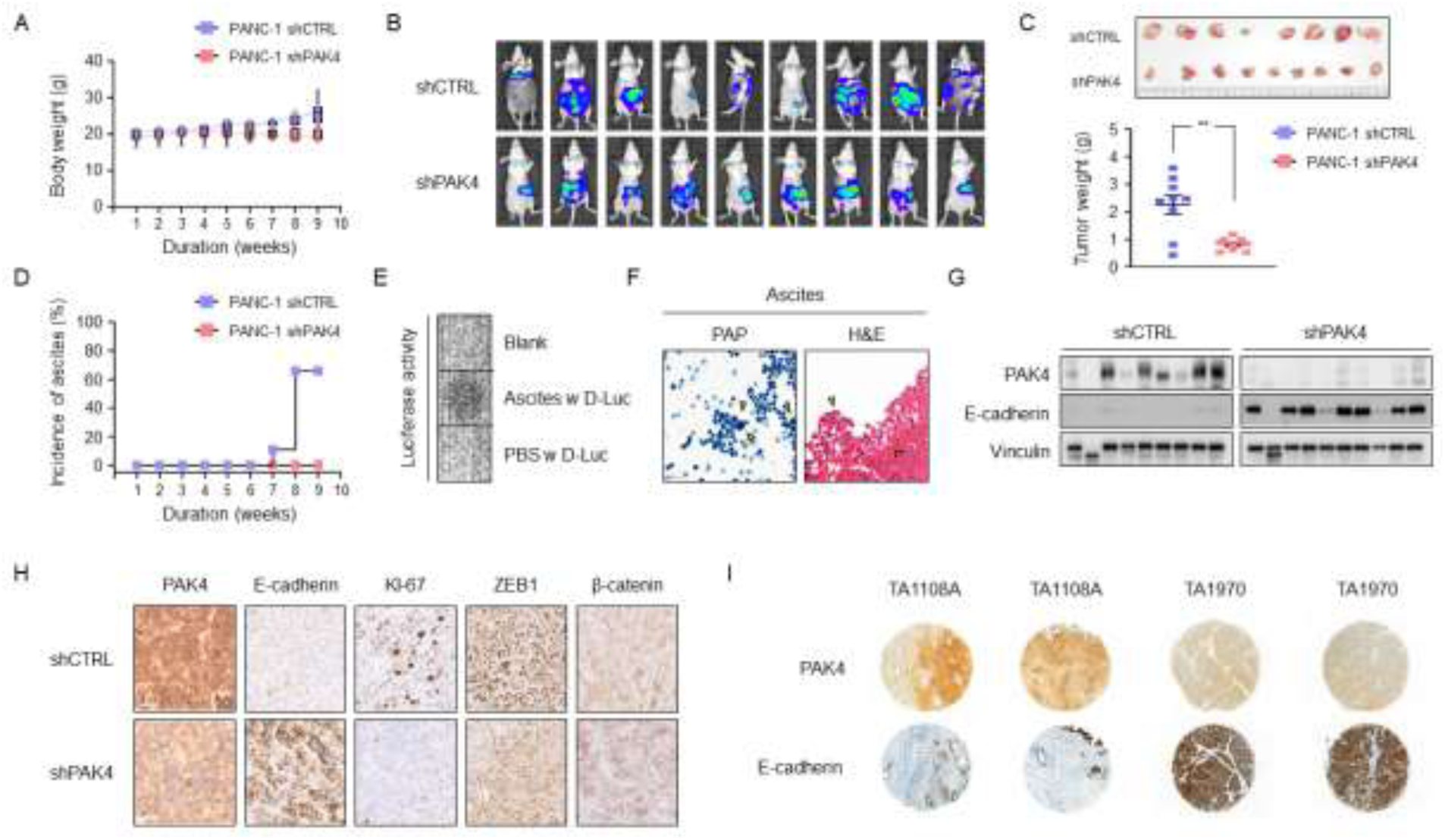
Knockdown of PAK4 inhibits primary pancreatic tumor growth, metastases, and incidence of ascites in an orthotopic pancreatic cancer mouse model. (A) The body weights of PANC-1 shCTRL and shPAK4 mice were measured over a period of 9 weeks. Values are presented as the mean ± standard error of the mean. A Student’s t-test was used to compare two independent groups. NS = no statistically significant difference. (B) Live animal imaging of mice at 9 weeks following orthotopic injection of PANC-1 shCTRL and shPAK4 cells. Primary tumors in the pancreas and metastases are visible. (C) Pancreatic tumors were dissected, weighed, and imaged (**, *P* < 0.01; *n* = 9, mean ± SEM). (D) Ascites were harvested using the peritoneal lavage method and (E) the presence of PANC-1 cells was confirmed with a luciferase assay. (F) Papanicolaou (PAP) and H&E staining were used to identify PANC-1 cells in ascitic fluid. (G) Western blot analysis and comparison of PAK4 and E-cadherin in primary pancreatic tumors from two different orthotopic mice. (H) Pancreatic tumor sections from PANC-1 shCTRL and shPAK4 mice were immunostained using PAK4, β-catenin, ZEB1, E-cadherin, and Ki-67 antibodies. (I) Immunohistochemistry for PAK4 and E-cadherin was performed on a tissue microarray that was produced using resected specimens from PDAC patients.

Consistent with our *in vitro* results, Western blot analysis and immunohistochemistry (IHC) staining of the primary tumors revealed that PAK4 knockdown significantly increased the expression of E-cadherin in PANC-1 cells compared to the control group (Fig 8G and 8H), while the expression of ZEB1 and Ki-67 was consistently decreased in the PAK4 knockdown tumors. However, the expression of β-catenin showed inconsistent changes upon PAK4 knockdown.

Finally, we performed IHC staining of resected specimens from patients with pancreatic cacner(*26*). For specimens with PAK4 overexpression, we observed a low number of E-cadherin positive cells. In contrast, some specimens with low expression of PAK4 exhibited a strong increase in E-cadherin positive cells compared to the PAK4 overexpression group (Fig. 8I). These data indicate that PAK4 and E-cadherin expression are regulated in a reciprocal manner in PDAC and support the hypothesis that inhibition of PAK4 may increase the expression of E-cadherin, subsequently leading to decreased migrasion, invasion and metastasis in PDAC patients.

## Discussion

In this study, we investigated the molecular mechanisms by which PAK4 overexpression contributes to disease progression and metastasis in PDAC. We found that PDAC cells harboring PAK4 overexpression displayed significantly higher levels of cell migration, invasion, and displacement compared with controls, suggestive of suppression of E-cadherin. By knocking down PAK4 in these cells, we observed stabilization of E-cadherin levels, which were modulated by the activity of Cdc42 and Hakai. Driving increased expression of PAK4 in PDAC cells with wild-type levels of PAK4 induced an increase in cell migration, invasion, and displacement, which linked to suppression of E-cadherin and elevated Cdc42 activity. These in vitro findings were replicated in an orthotopic mouse model. To the best of our knowledge, this is the first report demonstrating oncogene-like functions of PAK4 in PDAC cells, which directly interacts with E-cadherin and Hakai, as well as regulates the activity of Cdc42 to promote cell migration, invasion, and displacement.

To understand the association between PAK4 overexpression and cell migration in PDAC cells, we knocked down PAK4 in PDAC cells harboring PAK4 overexpression and evaluated whether silencing PAK4 alters cell migration. We observed that PAK4 knockdown dramatically inhibited single cell migration, collective cell migration, invasion, and cell displacement compared to control PDAC cells with PAK4 overexpression. In addition, suppression of PAK4 through knockdown modified sphere formation, reducing the irregular budding pattern observed in the control group. Several studies have suggested that EMT is involved in the early stages of cancer cell dissemination and is a critical factor in the incidence of invasion and metastasis in PDAC(*27, 28*). Interestingly, PAK4 knockdown in SU.86.86 and PANC-1 cells resulted in a striking upregulation of E-cadherin and alterations in a host of other EMT-related proteins, including ZEB1. These results suggest that PAK4 inhibition may repress EMT in PDAC cells, thereby altering their migratory and invasive potential.

ZEB1 is a transcription factor implicated in regulating EMT. It binds to specific DNA sequences, E-boxes, and the promoter region of the E-cadherin gene(*29, 30*), which leads to the recruitment of the chromatin-remodeling protein BRG1(*19*). BRG1 then alters the chromatin structure around the promoter region, making it less accessible to the transcriptional machinery and ultimately leading to a decrease in E-cadherin expression and induction of EMT. Similarly, β-catenin has been shown to play a significant role in the induction of EMT through a synergistic effect with TGF-β1 and cell adhesions(*31*). It is also involved in regulating the expression of E-cadherin through interactions at cell junctions(*32*). Thus, ZEB1 and/or β-catenin are capable of triggering EMT through the loss of cell-cell adhesion. Here, we found that PAK4 knockdown slightly inhibited the expression of ZEB1 in PDAC cells harboring PAK4 overexpression, but did not affect the nuclear translocation of β-catenin. In addition, we showed that PAK4 knockdown significantly attenuated the ability of TGF-β1 to increase SMAD signaling in PDAC cells, indicating that PAK4 plays a role in regulating E-cadherin stability through the non-canonical pathway undergoing EMT.

Cdc42 is a small GTPase that regulates cytoskeletal dynamics(*33*), localizes to cell-cell junctions, and contributes to the establishment and maintenance of cell polarity. Cdc42 has been shown to regulate EMT by affecting the stability of E-cadherin(*23*). In addition, bi-directional communication between E-cadherin and Cdc42 may be involved in the formation of filopodia(*34*). In the present study, when PAK4 was knocked down in PANC-1 cells, the stability of E-cadherin increased while the activity of Cdc42 decreased. Conversely, when PAK4 was overexpressed in AsPC-1 cells, E-cadherin stability decreased while the activity of Cdc42 increased. As shown with immunoprecipitation assays and immunofluorescence analysis, we discovered that PAK4 interacts with both E-cadherin, p120ctn, Cdc42 and colocalizes with these proteins. Furthermore, our study uncovered Hakai as a potential downstream target of PAK4 in the regulation of E-cadherin stability in PDAC.

To evaluate the metastatic potential of shCTRL and PAK4 knockdown in PANC-1 cells, we initially used a mouse xenograft model with cancer cells injected into the subcutaneous thigh (data not shown). However, we observed no difference in primary tumor growth and metastasis between the two groups: the shCTRL and the shPAK4 group. In contrast, in an orthotopic pancreatic cancer mouse model, PAK4 knockdown produced an anti-tumor effect, as demonstrated by decreased Ki-67 expression and increased TUNEL staining. Moreover, PAK4 knockdown reduced the incidence of malignant ascites and metastases compared to the shCTRL, without a significant change in body weight. The orthotopic model should therefore be suitable for evaluating metastatic potential.

In summary, our findings suggest that PAK4 plays the crucial role in regulating the cell signaling pathways that control the stability of E-cadherin, p120ctn, the activity of Cdc42, and Hakai. As a result, overexpression of PAK4 drives increased primary tumor growth as well as metastasis of PDAC cells. These findings provide insight into a potential role for PAK4 in PDAC progression and metastasis and suggest that targeting PAK4 could be a potential therapeutic strategy for PDAC patients with PAK4 overexpression.

## Materials and Methods

### In silico validation using the TCGA dataset

Survival data for PDAC patients (TCGA, Firehose Legacy, n = 181) with PAK4 wild-type or PAK4 overexpression were obtained from cBioPortal. For the TCGA dataset from cBioPortal.org(*35*), survival outcomes were calculated using the Kaplan-Meier method.

### In vitro cell culture

SNU-410, SNU-213, Capan-1, Capan-2, and ASPC-1 cells were obtained from the Korean Cell Line Bank (KCLB, Seoul, Republic of Korea) and cultured in RPMI-1640 (Welgene, catalog LM011-51) supplemented with 10% FBS (Gibco, catalog 26140079), 4mM L-glutamine (Welgene, catalog LM007-01), and 1% penicillin/streptomycin (Welgene, catalog LS202-2). MIAPaCa-2 cells were obtained from American Type Culture Collection (ATCC, CRL-1420) and grown in DMEM (Gibco, catalog 11995073) with 10% FBS, 2.5% horse serum (HS) (Gibco, catalog 26050088), 4mM L-glutamine, and 1% penicillin/streptomycin. PANC-1 and SU.86.86 cells were also obtained from ATCC (CRL-14369 and CRL1837, respectively) and grown in DMEM with 10% FBS, 4mM L-glutamine, and 1% penicillin/streptomycin. PaTu-8988T and PaTu-8988S were obtained from the DSMZ (Deutsche Sammlung von Mikroorganismen und Zellkulturen) and were grown in DMEM with 5% FBS and 5% HS, 4mM L-glutamine, and 1% penicillin/streptomycin. Cultures were maintained in a 5% CO_2_ incubator at 37°C. The authentication of each cell line was performed using the AmpFLSTR Identifiler PCR Amplification Kit (Applied Biosystems, catalog 4322288; Foster, CA, USA) by Macrogen (August 08, 2021). The 3530xL DNA Analyzer (Applied Biosystems) and the GeneMapper v5.0 software (Applied Biosystems) were used for DNA fingerprinting analysis.

### Cell proliferation assay

The cell proliferation assay was performed using the CellTiter-Glo Luminescent Cell Viability Assay (Promega, Madison, WI, USA) according to the manufacturer’s instructions. On day 0, 96-well plates were seeded with 2,000 cells/well. On day 3, plates were incubated for 1 h at room temperature, and 100 μL of CellTiter-Glo reagent was added to each well, followed by mixing on an orbital shaker for 10 min. Luminescence was measured on a Synergy H1 Hybrid Multi-Mode Reader from Bio Tek (Winooski, VT, USA).

### Fluorescence in situ hybridization analysis of PAK4 amplification

Fluorescence in situ hybridization analysis (FISH) was conducted as described previously (*36*). Briefly, a dual fluorescence kit (PathoVysion, Vysis, Abbott AG, Diagnostic Division Baar, Switzerland) was used to analyze the PAK4 gene on 2-µm thick paraffin sections. The probes were hybridized for 14–20 h at 37° C, washed twice in rapid wash solutions according to the manufacturer’s instructions, dried, and counterstained using DAPI. Slides were analyzed with an Olympus computer-guided fluorescence microscope.

### Plasmid, shRNAs, virus production, and transduction

pLKO.1 puro (Addgene #8453), pMDLg/pRRE (Addgene #12251), RSV/REV (Addgene plasmid #12253), pMD2.G (Addgene #12259), and pUMVC (Addgene #8449) were a gift from Prof. Bob Weinberg and Prof. Didier Trono(*37, 38*). pLenti CMV/TO V5-LUC Puro (Addgene plasmid #19785) was a gift from Prof. Eric Campeau and Paul Kaufman(*39*). pWZL Neo Myr Flag PAK4 was a gift from Prof. William Hahn and Prof. Jean Zhao(*40*). The sequence of interest was subcloned into the pLKO-puro backbone plasmids after digestion with AgeI/EcoRI (Table 1). The final shRNA constructs were confirmed with sequencing. Recombinant lentiviruses or retroviruses were produced by transfecting HEK 293T cells, using the Fugene6 Transfection Reagent (Promega, catalog E2691), with pCMV-VSV-G / pMD2.G (envelope plasmid), psPAX2 (lentivirus packaging plasmid) / pUMVC (retrovirus packaging plasmid), and lentiviral / retroviral constructs, according to the manufacturer’s instructions.

**Table 1.**
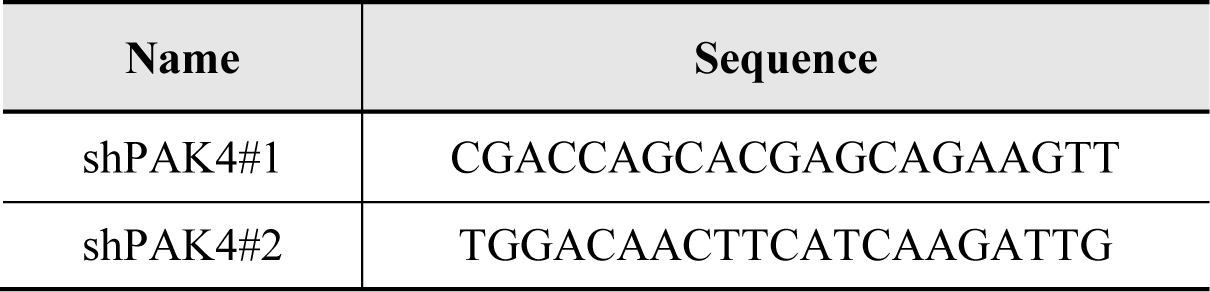
The sequence used for constructing shRNA plasmids.

### Single cell migration and invasion assay

Boyden chamber assays were performed according to the manufacturer’s instructions with 24-well inserts, uncoated for migration (Falcon, catalog 353097) or Matrigel-coated for invasion (Corning, catalog 354480). Cells were seeded in the upper chambers of a 24-well plate with serum-free medium. The lower chamber was filled with growth medium containing 10 ng/mL human recombinant TGF-β1 and 10 μg/mL mitomycin C. The 24-well plate was cultured at 37°C in a 5% CO_2_ humidified incubator for 20 hours, and cells remaining in the upper chamber above the filter membrane were removed with cotton swaps. Cells on the bottom of the membrane that had infiltrated the lower chamber were fixed with methanol and stained with 10 μg/mL Hoechst 33342 for 30 minutes. Finally, the stained cells were counted using an inverted fluorescence microscope (Zeiss Axio observer 7, Carl Zeiss, Germany). The average number of cells that had passed through the basement membrane in three random fields was used as the measure of cell invasiveness.

### Collective cell migration assay

Cells were seeded onto cover slips in 6-well plates and incubated for 24 hours. Subsequently, the cover slips were gently scratched using a 200 µL sterile tip. The cells at the partially detached edges of the scratch were allowed to reattach for 3 hours. Following this, microscope images were captured, defining the initial region (0 hour) using IncuCyte Zoom real-time imaging microscope (Essen Bioscience, Michigan, USA). The cells were then incubated for 24 hours to allow migration, with images taken at 12 and 24 hours.

### Spheroid forming assa

Cells were seeded at 1 × 10^4^ cells and incubated in 1% methylcellulose on poly–HEMA coated 24-well plates in RPMI1640 supplemented with 10% FBS, 1% PS, 20 ng/ml hFGF, 20 ng/ml hEGF, and B27. Suspension cultures were cultured for 14 days. Then, microscope images were taken by Zeiss Axio observer 7 (Zeiss, Jena, Germany).

### Cell displacement analysis

Cells were seeded in 96-well plates (ImageLock 4379, Sarorius Tokyo, Japan) and incubated for 24 hours. To observe cell polarity of shCTRL and shPAK4 PANC-1 cells, IncuCyte Zoom real-time imaging microscope (Essen Bioscience) was used to image cells every 15 minutes for 24 hours. Cell displacement was analyzeed by using the TrackMate plug-in in the ImageJ image processing program (*41*).

### Quantitative RT-PCR

Total RNA was prepared using TRIzol reagent (Invitrogen, Carlsbad, CA, USA) from cell lines according to the manufacturer’s recommendations. Synthesis of cDNA was performed using a PrimeScript™ RT reagent kit (TaKaRa Bio; Tokyo, Japan), and PCR was performed with Taq DNA Polymerase (Toyobo; Osaka, Japan). Real-time PCR was performed on the Thermal Cycler Dice® Real Time System III instrument (TaKaRa Bio; Tokyo, Japan), The reaction was performed as follows: 95°C for 30 seconds, followed by 40 cycles of 95°C for 5 secondss and then 60°C for 20 seconds. The primer sequences are listed in Table 2. 18s was used as the internal control

**Table 2.**
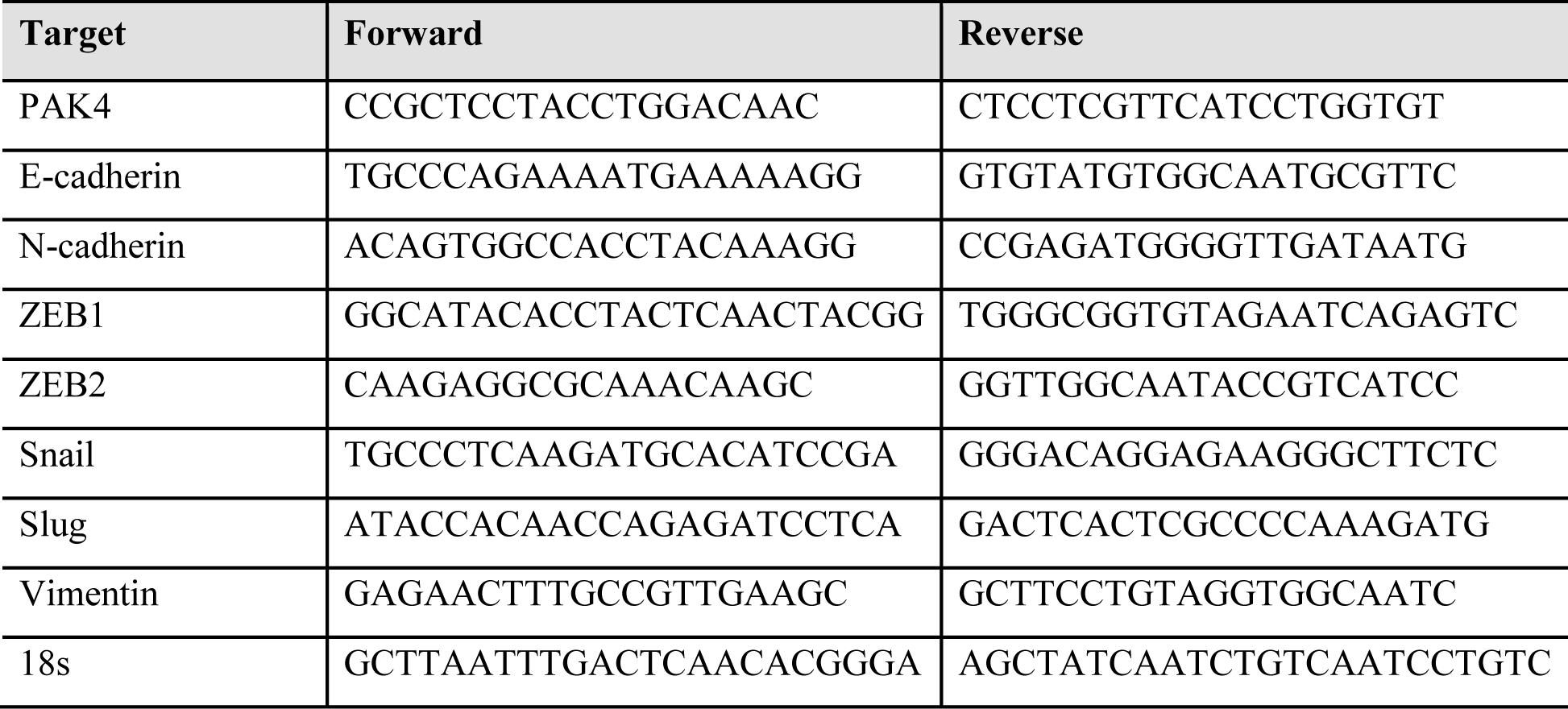
The primer sequences for qRT-PCR.

### Western blotting

Cell lysates were obtained by centrifugation at 12,000 × g for 30 minutes at 4°C. Protein concentration in the supernatant was measured by Bradford assay (BioLegend, San Diego, CA, USA). Proteins (10-40 μg) were separated by SDS polyacrylamide gel electrophoresis, and transferred to a polyvinylidene difluoride membrane (Bio-Rad, Hercules, CA, USA) that was blocked in a blocking buffer containing 5% skim milk, and then probed overnight with primary antibodies. Secondary antibodies conjugated with horseradish peroxidase (1:4,000 dilution; Bio-Rad) were applied for 1 hour. Immunoreactivity was detected by enhanced chemiluminescence (Biosesang, Seongnam, Republic of Korea) with a ChemiDoc Touch imager (Bio-Rad).

### Immunoprecipitation (IP) analysis

The cells were lysed with RIPA lysis buffer, and the whole cell lysates were incubated with appropriate antibodies at 4°C for 3 hours. Samples were then incubated for another 3 hours with protein A/G Sepharose beads. After washing the samples with RIPA lysis buffer, the immunoprecipitates were resolved on SDS-PAGE and probed with antibodies as indicated.

### Cdc42 and Rac1 activity assay

Cdc42 and Rac1 activity were evaluated using the Rac1/Cdc42 Activation Assay Kit (#17-441, Sigma) in accordance with the manufacturer’s instructions. Cells were grown in serum-free media for 20 hours and then lysed by scraping in Mg^2+^ lysis buffer. Following a brief centrifugation, the supernatant was collected. For every 400 μL of cell extract, 10 µg of Cdc42/Rac1 assay reagent (agarose beads) was added and incubated for 1 hour at 4°C with gentle rotation. Subsequently, beads were centrifuged (5 seconds, 13,000 rpm, 4°C), washed three times with wash buffer, and then resuspended in 4X SDS buffer, followed by boiling for 5 minutes at 95°C prior to western blotting. Total and activated Cdc42 and Rac1 were analyzed via western blotting, as previously outlined, using anti-Cdc42 or anti-Rac1 antibodies, respectively.

### Immunofluorescence microscopy

Cells were plated on coverslips, fixed with methanol for 1 h and blocked with 5 % bovine serum albumin in PBS for 1 h at room temperature, followed by incubation with primary antibodies at 4°C overnight. After 3 washes in PBS, the coverslips were incubated with secondary antibodies for 1 h at room temperature. All images were captured on Zeiss LSM880 confocal microscope (Carl Zeiss, Jena, Germany), with identical exposure times in each group.

### Orthotopic mouse model

All mice were housed in a specific pathogen-free facility at the Seoul National University Bundang Hospital, Seongnam, Republic of Korea. The project was approved by the Institutional Animal Care and Use Committee of Seoul National University Bundang Hospital (IACUC approval number: BA-2105-320-049-01). For orthotopic mouse studies, female athymic nude mice, weighing 20–22 g (5 weeks old), were purchased from Orient Bio Co. (Kapyong, Republic of Korea). Mice were provided with NIH-07 rodent chow (Zeigler Brothers, Gardners, PA, USA) purchased from Central Lab Animal Inc. (Seoul, Republic of Korea). Animals were acclimated to temperature (20–24 °C) and humidity (44.5–51.8%) and a 12 h light/dark cycle for one week before use. PANC-1 cells (1 × 10^7^) were implanted with Matrigel (BD Biosciences, San Jose, CA, USA) into the pancreas of each mouse.

The mice (n=9; shCTRL vs shPAK4 group) were weighed, and tumor areas and body temperature were measured throughout the study. The mice were maintained for eight weeks, and the mice were euthanized by CO_2_ asphyxiation, weighed, and subjected to necropsy. The volume and weights of xenograft tumors were recorded. Selected tissues were further examined by routine H&E staining and immunohistochemical analyses.

### In vivo optical imaging

Optical imaging was performed with a Xenogen IVIS200 small animal imaging system (Alameda, CA, USA) and was analyzed using Living Image software 4.3.1 (Caliper Life Sciences), as previously described(*42, 43*). Anesthesia (2.5% isoflurane) was administered in an induction chamber with 100% oxygen at a flow rate of 1 L/min and maintained in Xenogen IVIS 200 with a 1.5% mixture at 0.5 L/min. The athymic nude mice were injected with d-luciferin (100 mg/kg) dissolved in PBS (15 mg/mL) via the intraperitoneal route. Subsequently, the mice were placed in a prone position in the Xenogen IVIS200 and 1 to 5 min frames were consecutively acquired until the maximum signal was reached.

### Immunohistochemistry

Paraffin-embedded tissue blocks from the xenograft tumors were extracted and sectioned at a thickness of 5 μm. Tissue sections from mouse xenograft tumors, mounted on poly-L-lysine-coated slides, were deparaffinized by standard methods. Endogenous peroxidase was blocked by 3% hydrogen peroxide in PBS for 10 min. Antigen retrieval was done for 5 min in 10 mM sodium citrate buffer (pH 6.0) heated at 95°C in a steamer, followed by cooling for 15 min. Slides were washed with PBS and incubated for 1 h at room temperature with a protein-blocking solution (VECTASTAIN ABC kit, Vector Laboratories, Burlingame, CA, USA). Excess blocking solution was drained, and samples were incubated overnight at 4°C with one of the following: A 1:200 dilution of PAK4 (Abcam, ab62509), E-cadherin (Cell signaling, 3195), β-catenin (Cell signaling, 8480), zinc finger E-box binding homeobox 1 (ZEB1, Cell signaling, 3396), and Ki-67 (Invitrogen, MA5-14520) antibodies. Sections were then incubated with biotinylated secondary antibody followed by streptavidin (VECTASTAIN ABC kit). Color was developed by exposing the peroxidase to diaminobenzidine reagent (Vector Laboratories), which forms a brown reaction product. Sections were then counterstained with Gill’s hematoxylin (Sigma) for 1 min. Brown staining identified PAK4 (Abcam, ab62509), E-cadherin (Cell signaling, 3195), β-catenin(Cell signaling, 8480), ZEB1(Cell signaling, 3396), and KI-67 (Invitrogen, MA5-14520) expression.

### Ascites collection, papanicolaou (PAP), hematoxylin and eosin (H&E) staining

Ascitic fluid was collected from the abdominal region using peritoneal lavage. The smears stained with PAP were evaluated by Dr. Hee-Yong Na blinded to specimen information. The sediment smear method was used for the preparation of PAP staining. Unstained slides of the cells were stained with H&E.

#### TUNEL analysis

TUNEL apoptosis assay kit (Roche Diagnostics GmbH) was used to assess tumor apoptosis following fixation in 4% paraformaldehyde with vacuum overnight at room temperature. The representative slides were scanned into digital images with PANNORAMIC 250 Flash III (3DHISTECH, Budapest, Hungary). Images were annotated with CaseViewer (3DHISTECH, Budapest, Hungary).

### Statistical analysis

Statistical analyses were performed using SPSS v.12.0 software (SPSS Inc., Chicago, IL, USA). One-way analysis of variance was used for comparisons among groups. Significant differences between mean values were assessed with Duncan’s test. A p-value < 0.05 was considered statistically significant. The Student’s t-test was also used to compare two independent groups. *p < 0.05; **p < 0.01; or ***p < 0.001.

## Author Contributions

Conceptualization, K.J.K. and J.W.K.(Jin Won Kim); Methodology, K.J.K. and J.W.K.(Jin Won Kim); in vitro and in vivo experiments, K.J.K., J.H.S., B.R.P., and M.L.N.; software, K.J.K. and M.L.N.; validation, K.J.S., J.Y.L., J.O.L., S.H.K., J.W.K.(Ji-Won Kim), Y.J.K., K.W.L, J.H.K. S.M.B., and J.S.L.; formal analysis and investigation, K.J.K. and J.W.K.(Jin Won Kim); writing—original draft preparation, K.J.K., and J.W.K. (Jin Won Kim); writing—review and editing, K.J.K., J.M.S., E.H.J., S.A.K., M.S.K., K.J.S., J.Y.L., N.H.Y., J.W.K.(Ji-Won Kim), S.H.K., J.O.L., J.W.K.(Jin Won Kim), Y.J.K., K.W.L., J.H.K., Z.A.W., and S.M.B; visualization, K.J.K; supervision, J.W.K.(Jin Won Kim)

## Acknowledgments

This research was funded by Seoul National University Bundang Hospital Research Fund (No. 14-2023-0032) and the National Research Foundation of Korea (NRF) grant funded by the Korean government (MIST) (No. 2020R1F1A1076372 and RS-2023-00240907).

