## Supplementary material for "Loss of p21-activated kinase 4 (PAK4) suppresses pancreatic tumor progression and metastasis through regulating E-cadherin": Supplementary information_Uncropped images.pdf

Supplementary Figure 1. Uncropped western blots related to Figure 2A

PAK4

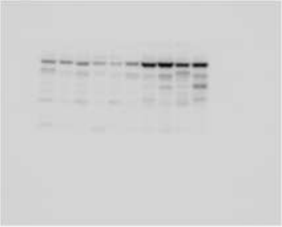

GAPDH

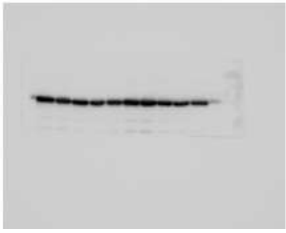

Supplementary Figure 2. Uncropped western blots related to related to Figure 3A

PAK4

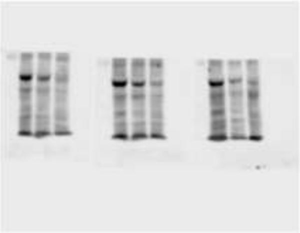

Vinculin

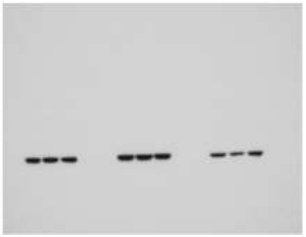

Supplementary Figure 3. Uncropped western blots related to related to Figure 4B

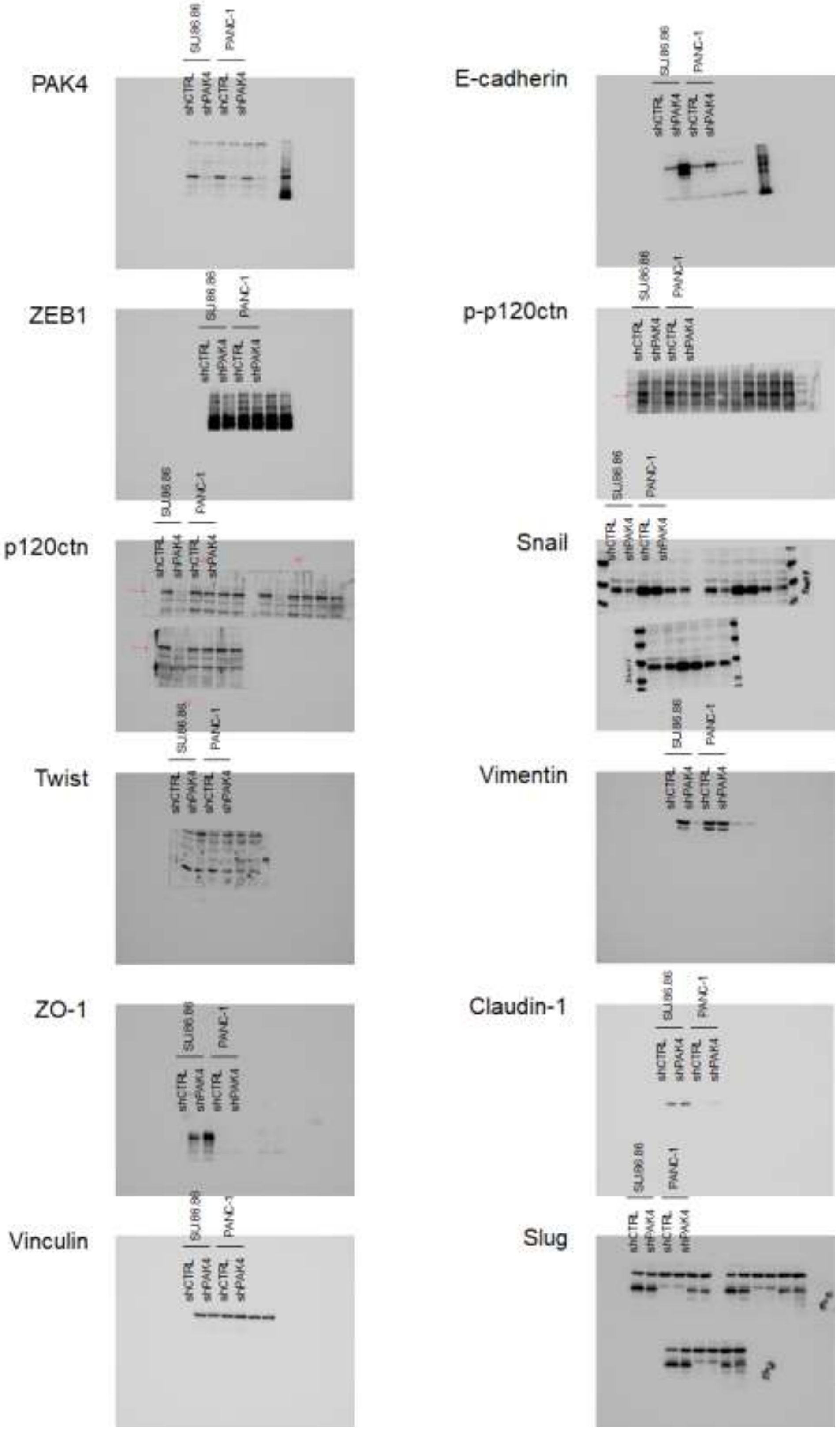

Supplementary Figure 4. Uncropped western blots related to related to Figure 4C

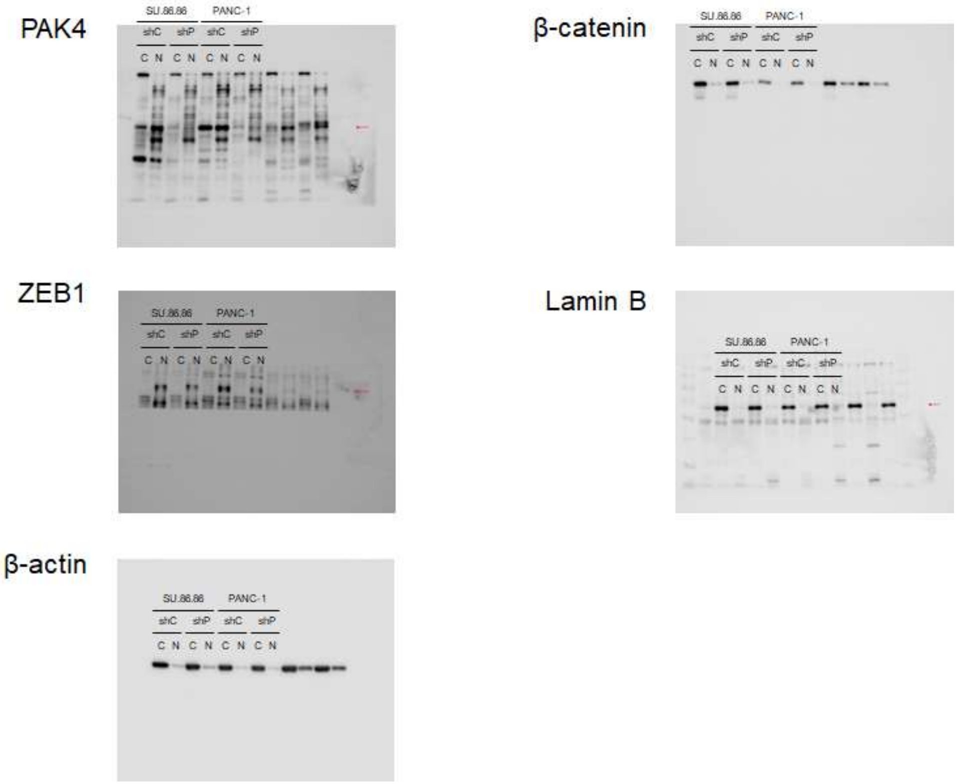

Supplementary Figure 5. Uncropped western blots related to related to Figure 4E

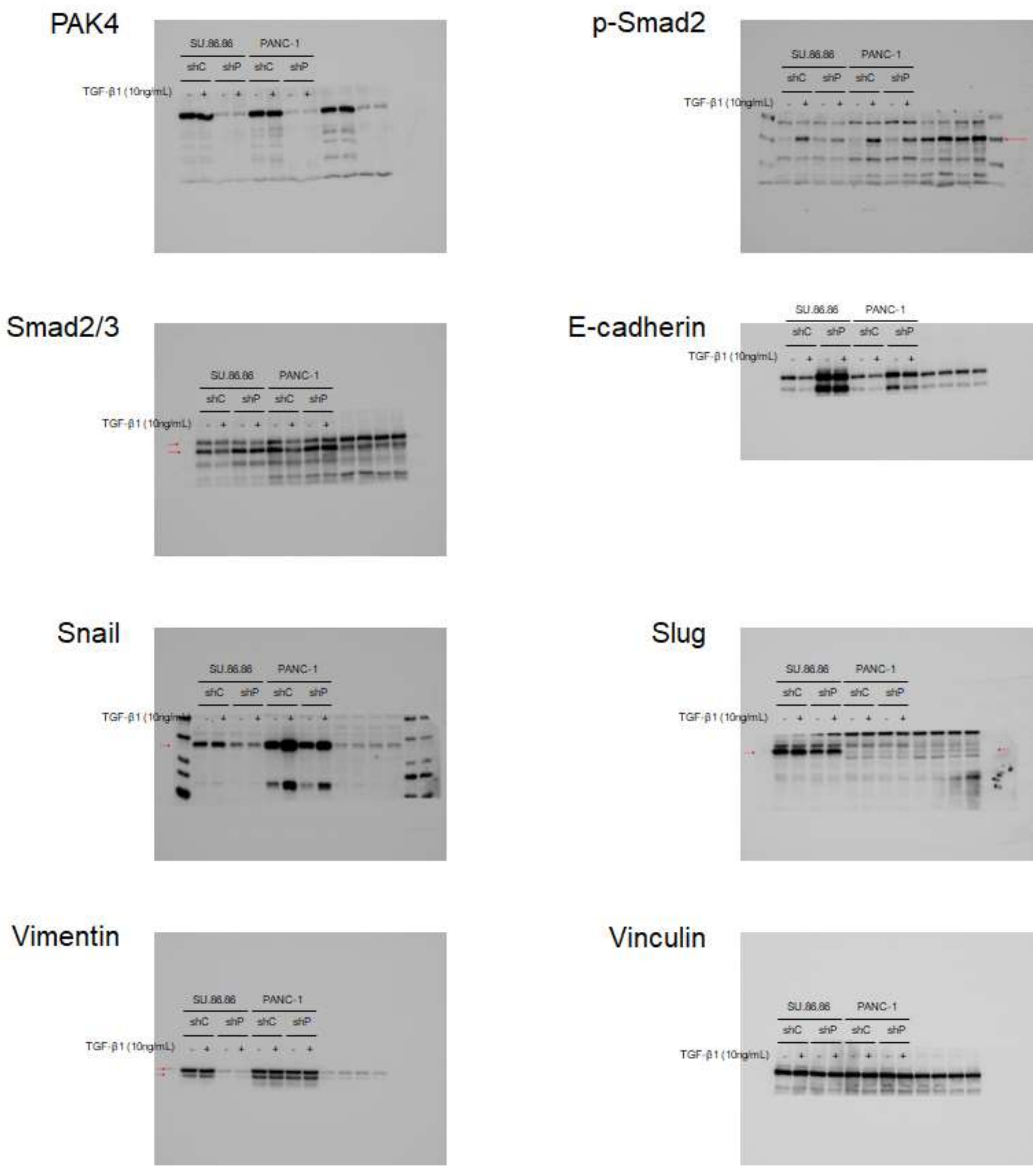

Supplementary Figure 6. Uncropped western blots related to related to Figure 5A

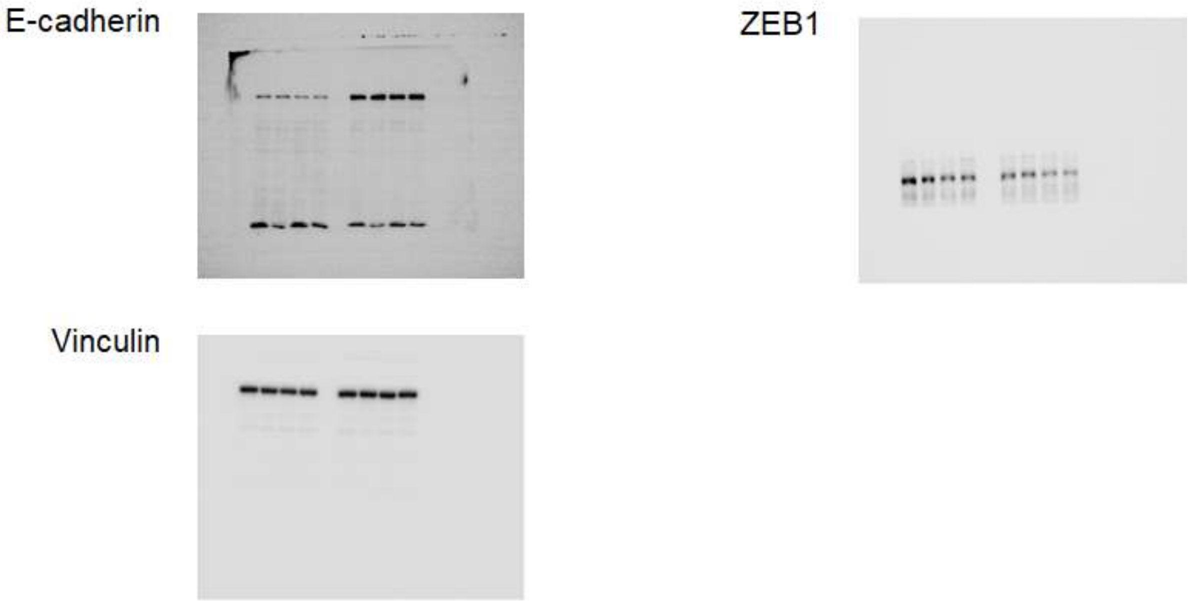


Supplementary Figure 7. Uncropped western blots related to related to Figure 5B

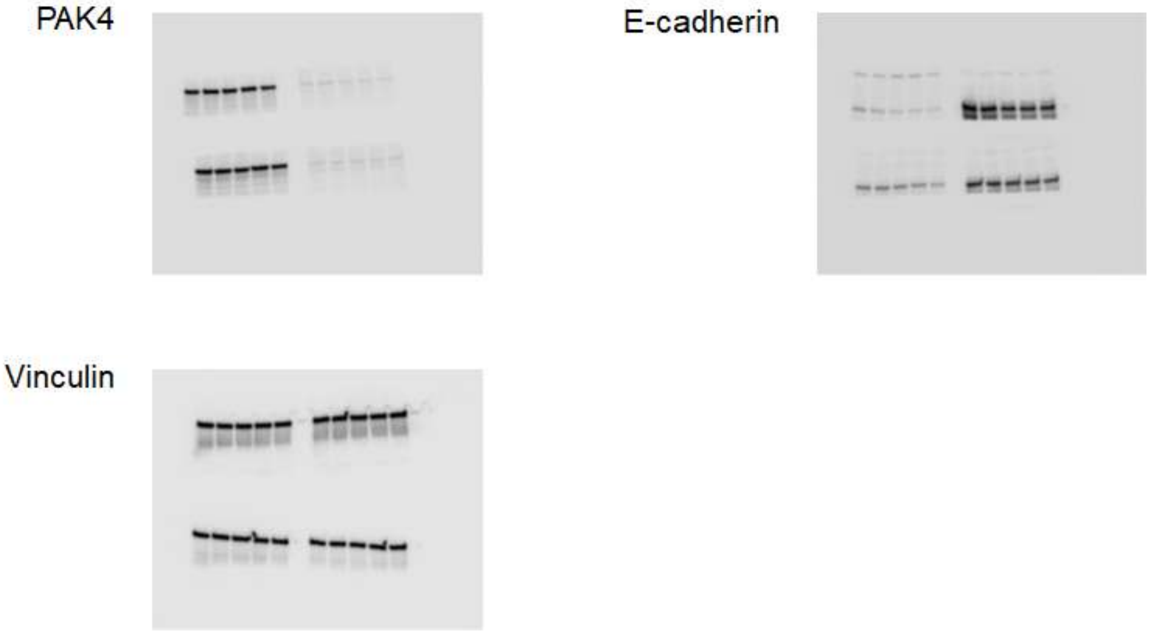

Supplementary Figure 8. Uncropped western blots related to related to Figure 5C

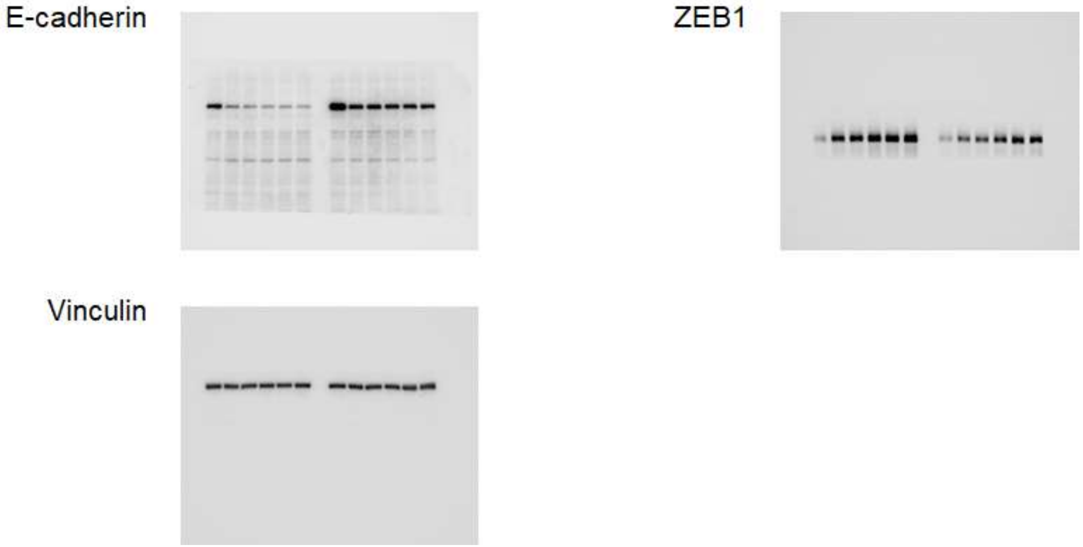


Supplementary Figure 9. Uncropped western blots related to related to Figure 5D and 5E

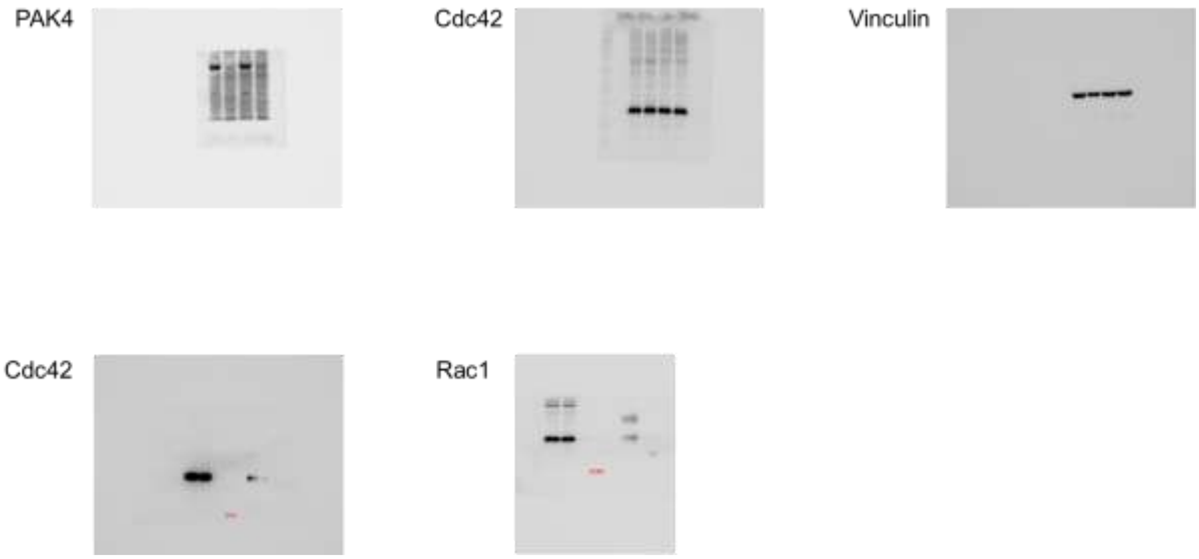

Supplementary Figure 10. Uncropped western blots related to related to Figure 5F and 5G

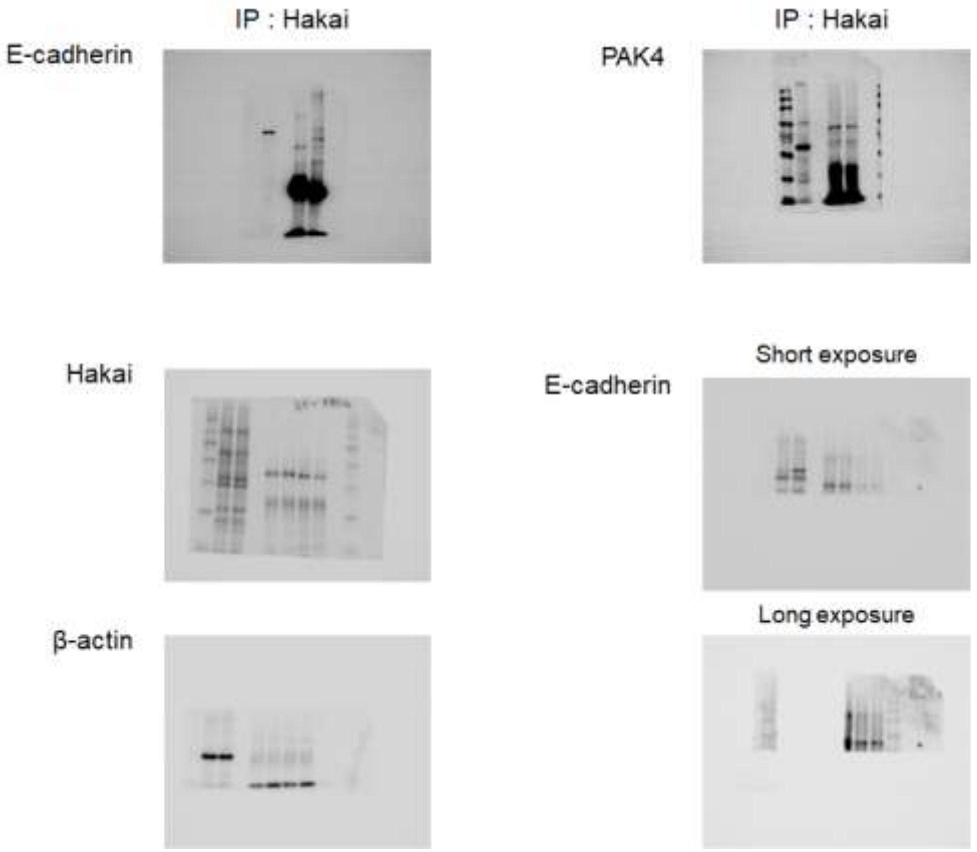

Supplementary Figure 11. Uncropped western blots related to related to Figure 6A

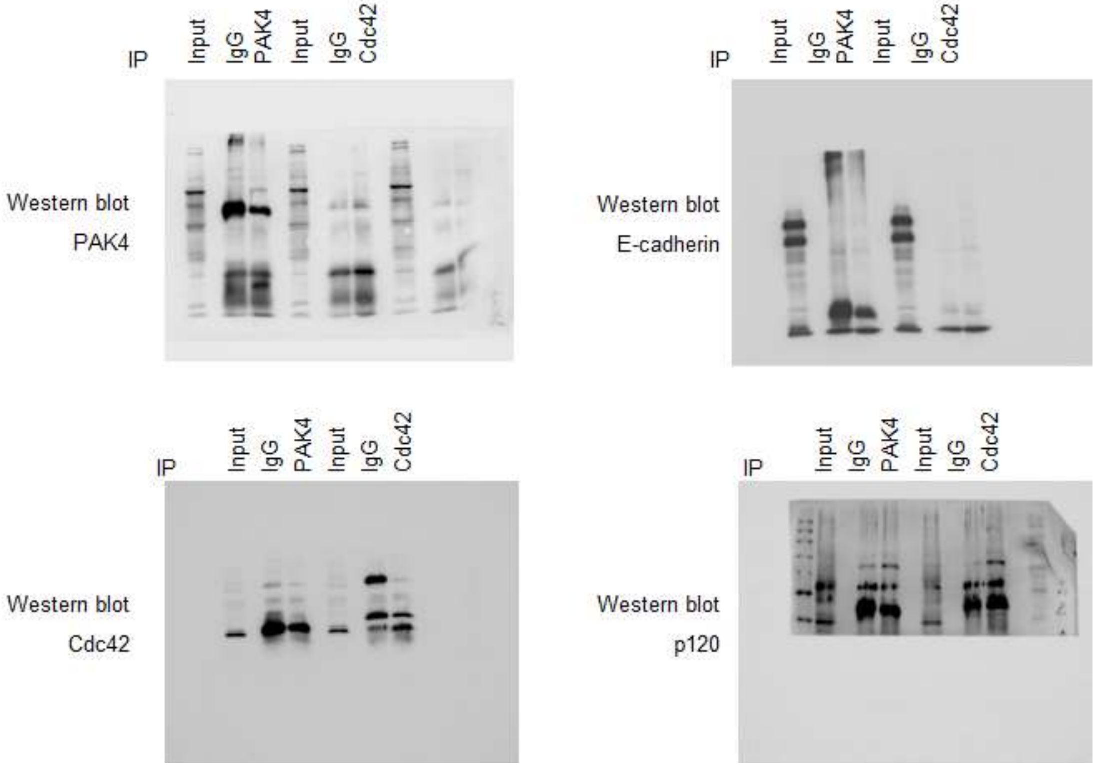

Supplementary Figure 12. Uncropped western blots related to related to Figure 7A and 7E

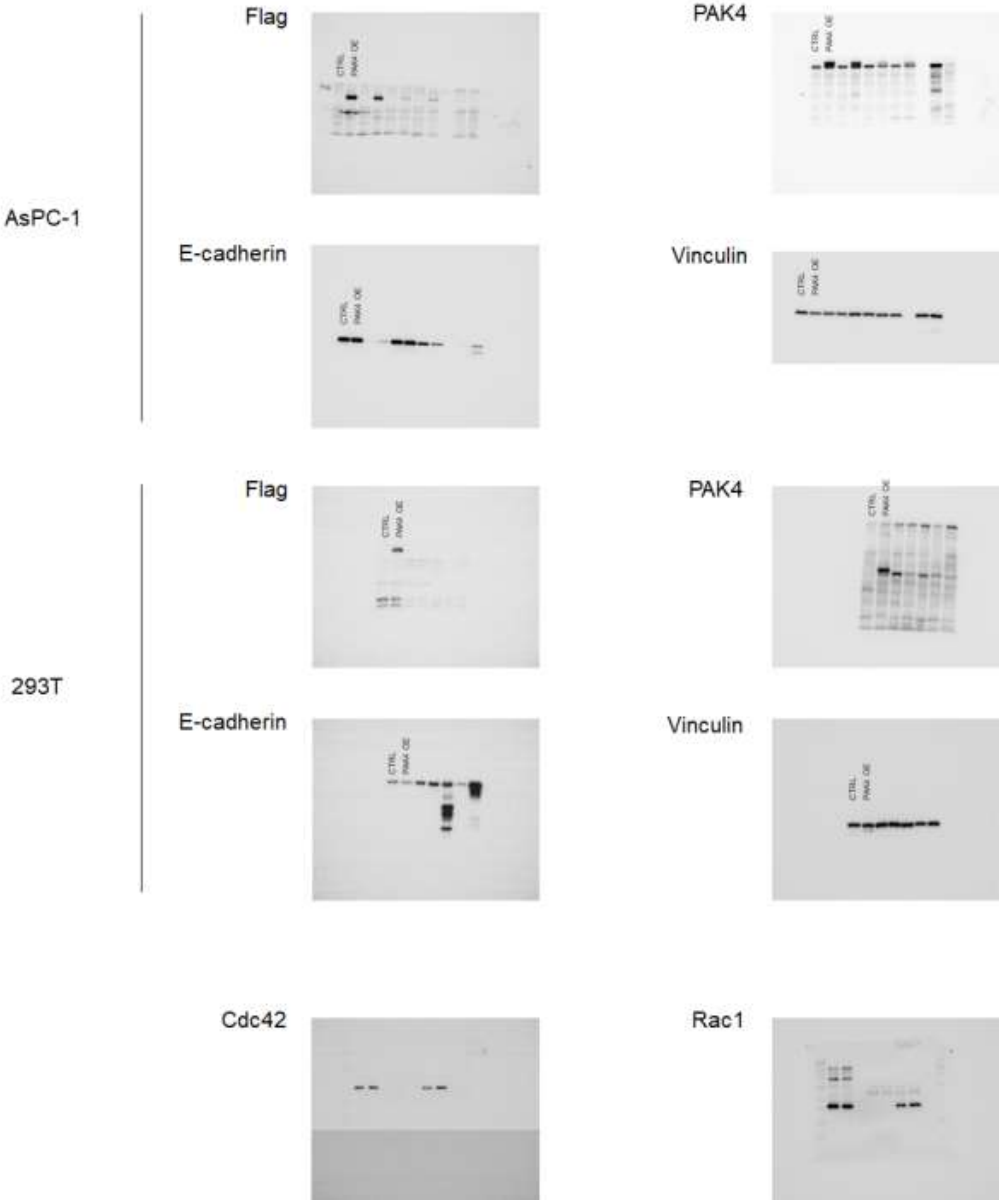

Supplementary Figure 13. Uncropped luciferase activity image related to related to Figure 8E

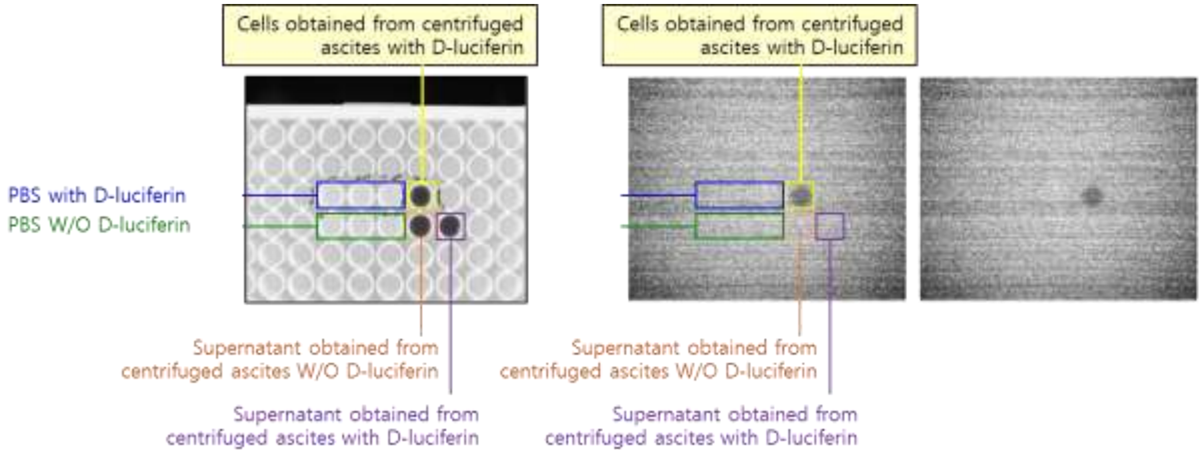


Supplementary Figure 14. Uncropped H&E and PAP staining images related to related to Figure 8F

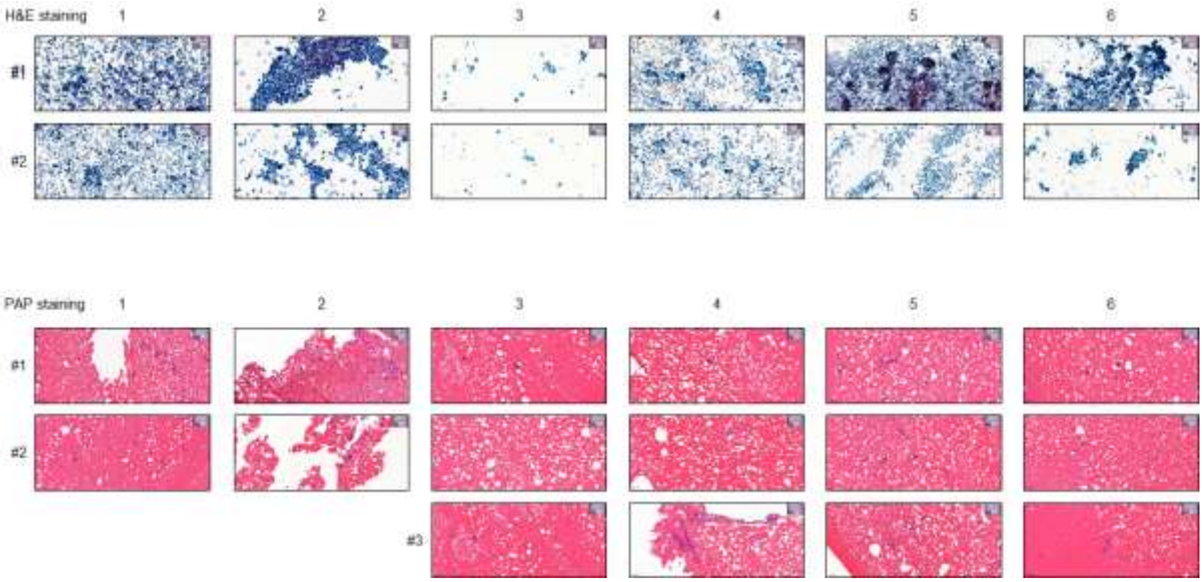

Supplementary Figure 15. Uncropped western blots related to related to Figure 8G

PAK4

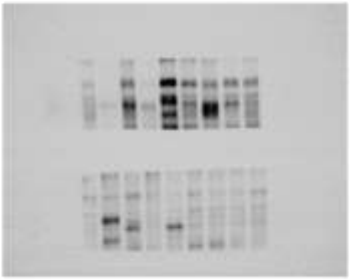

E-cadherin

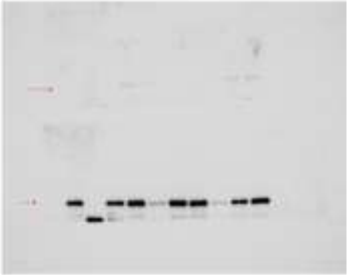

Vinculin

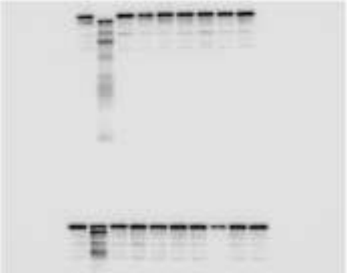
