## Supplementary figures and images for "Loss of p21-activated kinase 4 (PAK4) suppresses pancreatic tumor progression and metastasis through regulating E-cadherin"

### Graphical abstract.jpg

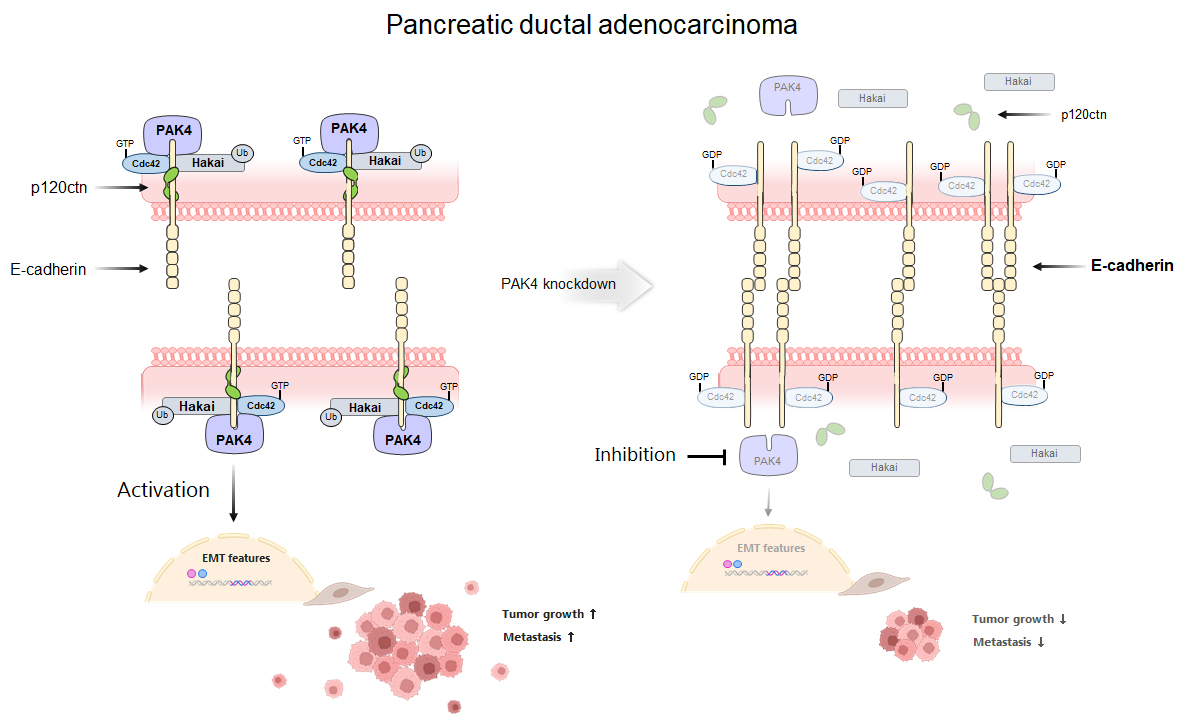
